# AMLs harboring DNMT3A-destabilizing variants show increased intratumor DNA methylation heterogeneity at bivalent chromatin domains

**DOI:** 10.1101/2023.02.13.528223

**Authors:** Dohoon Lee, Bonil Koo, Seok-Hyun Kim, Jamin Byun, Junshik Hong, Dong-Yeop Shin, Choong-Hyun Sun, Ji-Joon Song, Jaesung Kim, Siddhartha Jaiswal, Sung-Soo Yoon, Sun Kim, Youngil Koh

## Abstract

The mechanistic link between the complex mutational landscape of *de novo* methyltransferase *DNMT3A* and the pathology of acute myeloid leukemia (AML) has not been clearly elucidated so far. A recent discovery on the catalogue of DNMT3A-destabilizing mutations throughout the *DNMT3A* gene as well as the oligomerization-dependent catalytic property of DNMT3A prompted us to investigate the common effect of DNMT3A-destabilizing mutations (*DNMT3A*^INS^) on the genomewide methylation patterns of AML cells. In this study, we describe the characteristics of *DNMT3A*^INS^ AML methylomes through the comprehensive computational analyses on three independent AML cohorts. As a result, we show that methylomes of *DNMT3A*^INS^ AMLs are considerably different from those of *DNMT3A*^R882^ AMLs in that they exhibit both locally disordered DNA methylation states and increased across-cell DNA methylation heterogeneity in bivalent chromatin domains. This increased epigenetic heterogeneity was functionally associated with heterogeneous expression of membrane-associated factors shaping stem cell niche, implying the diversification of the modes of leukemic stem cell-niche interactions. We also present that the level of methylation disorder at bivalent domains predicts the response of AML cells to hypomethylating agents through cell line- and patient-level analyses, which supports that the survival of AML cells depends on stochastic DNA methylations at bivalent domains. Altogether, our work provides a novel mechanistic model suggesting the genomic origin of the aberrant epigenomic heterogeneity in disease conditions.

## Introduction

Recent sequencing efforts of acute myeloid leukemia (AML) genomes and exomes have identified *DNMT3A* as one of the most recurrently mutated epigenetic modifiers whose mutation is associated with adverse patient outcome^1^. *DNMT3A* encodes a *de novo* DNA methyltransferase that establishes DNA methylation patterns during the development of mammalian stem cells^2^, but the precise molecular mechanism underlying the initiation and progression of AML mediated by mutant DNMT3A has not been clearly elucidated. One of the characteristics that obscures the identification of the mechanistic role of mutant DNMT3A in AML is its intricate mutational landscape. In AML, about 60% of the *DNMT3A* mutations cause amino acid substitution of arginine at position 882 (R882) and the remaining ∼40% of mutations are seemingly dispersed throughout the functional domains of *DNMT3A*^3^. Thus, much attention so far has been primarily drawn on the significance of *DNMT3A* R882 mutations in AML due to their prevalence. The results of such studies are gradually reaching at the consensus that mutant DNMT3A^R88^^2^ elicits dominant negative effect by hampering wildtype DNMT3A from forming catalytically active homotetramers^4^, in spite of some opposing results^5^. On the contrary, for *DNMT3A* mutations other than the R882 mutation (non-R882 mutations), much of their clinical implication and mechanistic role in AML pathogenesis still remain to be elucidated. Recently, a comprehensive biochemical characterization of 253 variants across *DNMT3A* gene suggested that a considerable number of disease-associated *DNMT3A* variants trigger the destabilization of the protein followed by its proteasomal degradation^6^. Intriguingly, these variants inducing the instability of DNMT3A (*DNMT3A*^INS^), and perhaps reduced intracellular concentration of intact DNMT3A, seemed to confer high fitness advantages to the cells of hematopoietic lineage, but the underlying molecular mechanism linking *DNMT3A*^INS^ and the progression of hematological disorders has not been clarified thoroughly.

Meanwhile, the epigenetic diversity of cancer cells, primarily in terms of the heterogeneity of DNA methylation patterns, is increasingly acknowledged as an important factor that contributes to the increased adaptive potential of the tumor, which leads to adverse outcome, treatment resistance, or shorter interval to relapse rate in a variety of cancer types^7–9^. In chronic lymphocytic leukemia, it has been reported that locally disordered methylation patterns at promoter regions are associated with increased transcriptional variability as well as adverse patient outcomes^7^, and its implication for the treatment resistance and disease relapse has been reported in diffuse large B-cell lymphoma^10^. The role of DNA methylation heterogeneity in AML has also been studied recently^11^. Given these broad clinical implications of DNA methylation heterogeneity, it has been widely accepted that the increased fitness of cancer cell population conferred by the epigenetic diversity is pivotal. However, the connection between a specific subset of DNMT3A variants and the extent of disorder of DNA methylation patterns have not been characterized so far.

Here, we provide a molecular-level insight into the fitness advantages conferred by *DNMT3A*^INS^ variants through the investigation of their overall impact on the DNA methylomes and transcriptomes of AML patients. Particularly, we explore the association between *DNMT3A*^INS^ and the disorderedness of DNA methylation patterns, in addition to the DNA methylation features that are routinely analyzed, such as promoter methylation levels or differentially methylated regions (DMRs). For the direct and robust examination of the methylomes of AML patients with *DNMT3A*^INS^, we extensively reanalyzed publicly available methylation profiles of AML patients from the two large independent cohorts^8, 12^. Furthermore, we performed reduced-representation bisulfite sequencing (RRBS) on our own cohort for validation. Through these analyses on diverse cohorts, we show *DNMT3A*^INS^ AMLs exhibit increased local DNA methylation disorder as well as epigenetic cellular diversity that are associated with the transcriptional heterogeneity of genes having roles in determining the leukemic stem cell niche. Given the previous studies showing the oligomerization-dependent shift of catalytic processivity of DNMT3A and the concentration-dependent oligomerization preference of DNMT3A, this study suggests an interesting model of pathogenesis having *DNMT3A*^INS^ variants as the genetic origin of epigenetic instability.

## Results

### Definition of *DNMT3A*^INS^ variants

To obtain a predefined set of *DNMT3A*^INS^ variants, we utilized previous experimental results of the protein stability assay measuring the stability scores of mutant DNMT3A protein in terms of the stability ratio normalized to WT DNMT3A^6^. From the stability ratios for 253 disease-associated variants affecting 248 unique amino acid residues, we could obtain stability scores for each of the 248 residues by assigning average stability ratios for all substitutions associated with that residue. Since the resulting stability scores displayed a bimodal distribution, we could naturally divide them into two groups, namely destabilizing (n=125) and non-destabilizing (n=123) residues, based on the score 0.75 (Figure 1a, Supplementary Table 1). To further justify this grouping, we investigated the full-length structure of DNMT3A (obtained from AlphaFold Protein Structure Database^13^, Uniprot ID Q9Y6K1) and found that destabilizing residues are enriched in β-sheets behind the helical tetramer interface compared to non-destabilizing residues (Supplementary Figure 1a-d). Furthermore, destabilizing residues showed higher predicted local distance difference test (pLDDT) values, which generally represent greater evolutionary conservation and structural importance of the residues (Supplementary Figure 1e). Given these biochemical, structural and evolutionary grounds, we defined a *DNMT3A*^INS^ variant as a point mutation occurring at destabilizing residues as well as nonsense and frameshift mutations occurring at any position of the protein to cover a broader spectrum of instability-inducing variants. Meanwhile, point mutations occurring at non-destabilizing positions other than R882 were defined as *DNMT3A*^Other^ variants.

**Figure 1.**
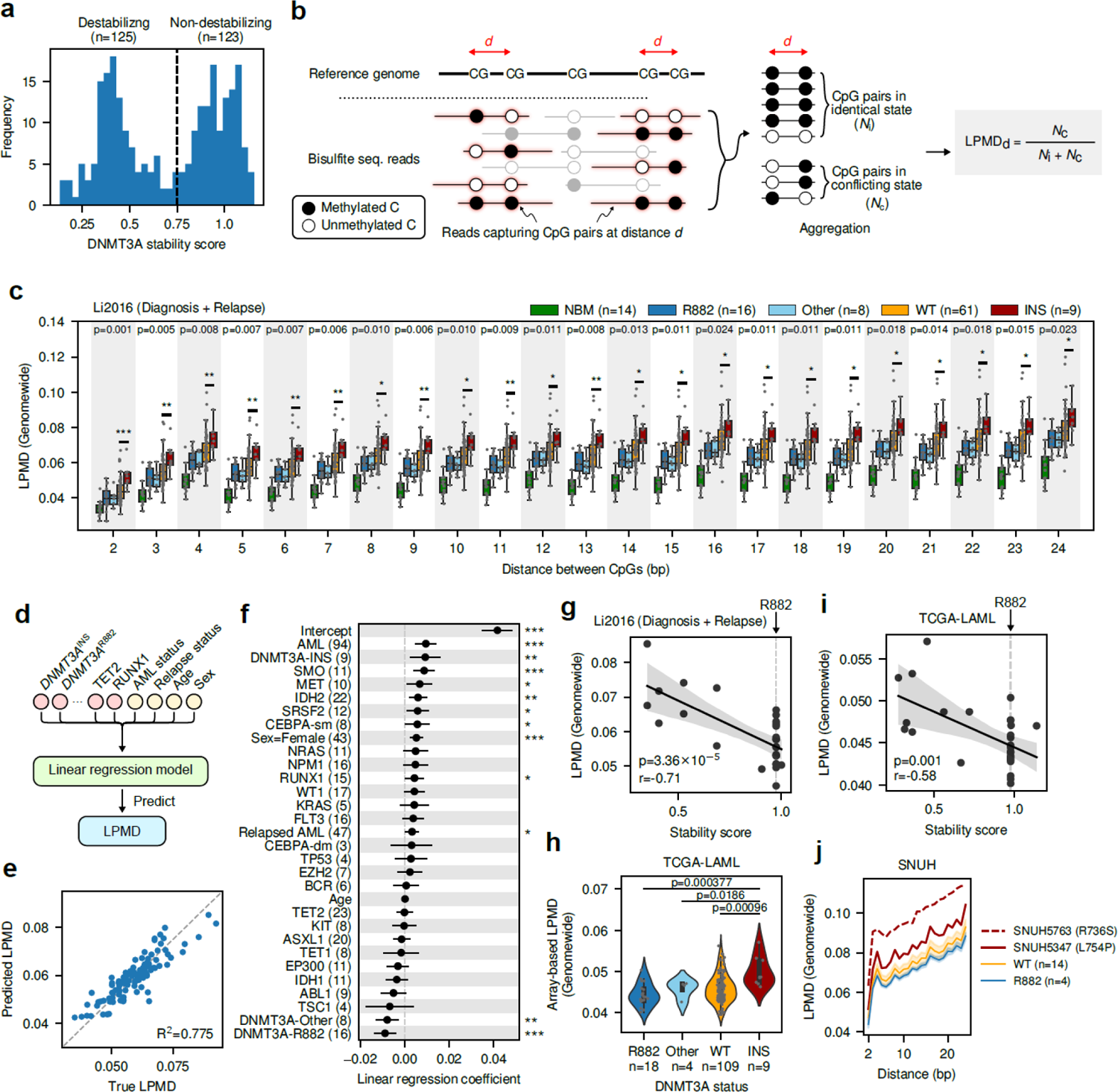
DNMT3AINS AMLs show locally disordered DNA methylation patterns. (a) Distribution of DNMT3A stability scores for 248 residues across DNMT3A protein. The dotted line denotes the threshold value 0.75 dividing destabilizing and non-destabilizing residues. (b) Description of local pairwise methylation discordance calculation. (c) Comparison of genomewide LPMD between different DNMT3A mutation subclasses in diagnosis and relapse AML samples from Li2016 cohort. P-values from two-sided Mann-Whitney U tests between DNMT3AWT and DNMT3AINS subclasses are shown. (d) Schematic diagram illustrating the multiple linear regression analysis predicting LPMD values based on mutation status, age and gender. (e) Accuracy of LPMD values predicted by multiple linear regression analysis. (f) Coefficients and significances of regression coefficients. (g) Correlation between DNMT3A stability score and genomewide LPMD in Li2016 cohort. Pearson’s correlation coefficient and corresponding p-value is shown. (h) Array-based LPMD of TCGA-LAML samples. P-values from two-sided Mann-Whitney U tests are shown. (i) Correlation between DNMT3A stability score and genomewide LPMD in TCGA-LAML cohort. Pearson’s correlation coefficient and corresponding p-value is shown. (j) Genomewide LPMD comparison in SNUH cohort. In (c), *** p < 0.001, ** p < 0.01, * p < 0.05, two-sided Mann-Whitney U test; The center line denotes the median, the upper and lower box limits denote upper and lower quartiles, and the whiskers denote 1.5× interquartile range. In (f), CEBPA-sm, CEBPA with single mutation; CEBPA-dm, CEBPA with double mutation.

### *DNMT3A*^INS^ AMLs show locally disordered DNA methylation patterns

DNMT3A exerts its catalytic activity by forming oligomers. Intriguingly, the mechanism of DNMT3A-mediated *de novo* methylation is shown to be dependent on its oligomeric state^14^. A homotetrameric complex exhibits processive catalysis in which the addition of methyl group occurs consecutively on CpGs within a local stretch of DNA, whereas a dimeric complex shows distributive catalysis in which the complex rapidly dissociates from the DNA after a catalysis. Since the oligomeric state of DNMT3A was shown to be dependent on the intracellular concentration of the protein^15^, we hypothesized that the distributive *de novo* methylation mediated by dimeric DNMT3A will be prevalent in *DNMT3A*^INS^ AMLs. To quantify the extent of the processive or distributive *de novo* methylation from the traces left on the methylomes of AML patients, we utilized a computational measure called local pairwise methylation discordance^16^ (LPMD; Figure 1b). LPMD is a per-sample measure that represents the extent to which a pair of nearby CpGs have different methylation states. Since the processive methylation will make a pair of CpG sites at a close distance both methylated, LPMD in turn reflects the processivity of *DNMT3A*, even though we cannot simply rule out the effects of other factors including TET-driven demethylation.

We conducted a reanalysis of the enhanced reduced-representation bisulfite sequencing (eRRBS) data provided by Li et al.^8^ (hereafter called Li2016 cohort) for 94 paired diagnosis and relapse samples from 47 AML patients. We first identified somatic mutations for all the 94 AML samples and compared their LPMDs altogether according to their *DNMT3A* mutation states. As expected, LPMD steadily increased as the distance between CpG pairs increased, reflecting the local homogeneity of DNA methylation states (Figure 1c). Surprisingly, we observed that *DNMT3A*^INS^ AMLs showed significantly higher genomewide LPMD than any other *DNMT3A* subclasses (p=0.001, two-sided Mann-Whitney U test between WT and *DNMT3A*^INS^ for 2bp-away CpG pairs; Figure 1c), suggesting the dysregulation of local correlations of DNA methylation states in *DNMT3A*^INS^. To ensure that the association between *DNMT3A*^INS^ and locally disordered methylation states remains significant even after accounting for other co-occurring mutations, ages, and genders, we built a multivariate linear regression model predicting LPMD (Figure 1d, e) and found that the association between *DNMT3A*^INS^ and high LPMD value was indeed significant after adjusting for such factors (Figure 1f). Notably, *DNMT3A*^INS^ was shown to be the only *DNMT3A* mutation subclass that was positively associated with LPMD (multiple linear regression coefficient of 0.0095), which was in stark contrast to the negative association of the other *DNMT3A* mutation subclasses (multiple linear regression coefficient of −0.0093 and −0.0083 for *DNMT3A*^R882^ and *DNMT3A*^Other^, respectively) on LPMD. It is worth noting that the age did not show significant correlation with LPMD values, suggesting that the contribution of aging-associated alterations of methylation patterns is insignificant in this case (Figure 1f). We verified that bisulfite conversion rates were greater than ∼99.7% for all the examined eRRBS data (median 99.87%) and also were not correlated with LPMD values, thus excluding the possibility that the high LPMD occurring due to experimental artifacts (Supplementary Figure 2a, b).

We next examined whether the extent of the destabilization of DNMT3A induced by *DNMT3A*^INS^ mutation correlates with LPMD. We found that the stability scores showed marked negative correlation with LPMD values (Pearson’s r=-0.71, p=3.36×10^-5^; Figure 1g). In other words, more severe instability of DNMT3A was associated with greater local discordance of DNA methylation states. This result corroborates the putative relationship between the instability-driven reduction of intracellular DNMT3A concentration and increased DNA methylation disorder.

To verify whether these findings can be reproduced in an independent AML cohort, we conducted similar analysis for the TCGA-LAML cohort (n=140). Since we only had methylation BeadChip array profiles for this cohort, we could not make use of the phasing information of methylation states as in the bisulfite sequencing data from Li2016 cohort. To circumvent this problem, we instead devised an array-based LPMD as an approximation of bisulfite sequencing-based LPMD (Methods) and computed it for the TCGA-LAML cohort. Of note, array-based LPMD serves as a lower bound of sequencing-based LPMD. As a result, we observed that *DNMT3A*^INS^ AMLs had significantly high levels of local disorder of DNA methylation (Figure 1h). Furthermore, the array-based LPMD levels were also negatively correlated with the stability scores of corresponding *DNMT3A* variants (Figure 1i; Pearson’s r=-0.58, p=0.001), reproducing the results from Li2016 cohort.

Additionally, we newly performed RRBS on our own cohort comprised of 20 AML patients (SNUH cohort; Supplementary Table 2). There were two patients with *DNMT3A*^INS^ variants at position 754 (stability score 0.386) and 736 (stability score 0.316). Of note, these variants were among the highly critical variants impacting the stability of the protein (top 17% and 7%, respectively). Again, those two *DNMT3A*^INS^ AML patients showed markedly high genomewide LPMD values (Figure 1j). We confirmed that variant at 736 position is provoking decreased tetramerization at protein level with prominent formation of dimerization (Supplementary Figure 3).

Given the difference of *DNMT3A*^INS^ and *DNMT3A*^R882^ in terms of local DNA methylation disorderedness, we then asked whether the difference of *DNMT3A*^INS^ and *DNMT3A*^R882^ AMLs can also be found in their mutational co-occurrence patterns. We conducted mutational co-occurrence analyses using TCGA-LAML (n=180), BeatAML (n=281) and Leucegene (n=263) cohorts. Although there was substantial inter-cohort difference of mutational co-occurrence and mutual exclusivity patterns (Supplementary Figure 4a-c), we observed that *DNMT3A*^INS^ and *DNMT3A*^R882^ AMLs did not share similar mutational patterns in both cohort-wise (Supplementary Figure 4a-c) and pooled (Supplementary Figure 4d) analyses except for the co-occurrence with *NPM1* mutations. These results, along with the remarkable difference in local disorder of DNA methylation between *DNMT3A*^INS^ and *DNMT3A*^R882^, prompted us to seek for a deeper understanding of the mechanistic difference between *DNMT3A*^INS^ and *DNMT3A*^R882^ AMLs in terms of their global methylation landscapes.

### Methylation landscape of *DNMT3A*^INS^ AMLs, in terms of methylation levels, is similar to that of *DNMT3A*^WT^ AMLs, but not *DNMT3A*^R882^ AMLs

In general, it is widely known that the alteration of DNA methylation in cancer cells accompanies focal hypermethylation of CpG-dense regulatory regions including CpG islands, as well as a global loss of DNA methylation. AML cells are no exception to these epigenomic alterations. Beyond these malignancy-associated alterations, *DNMT3A*^R882^ AMLs are shown to have distinct hypomethylation patterns compared to *DNMT3A*^WT^, which arise from the attenuated AML-associated hypermethylation and loss of methylation at regions normally maintained at high methylation level^17^. On the other hand, the characteristic of the global methylation landscape of *DNMT3A*^INS^ AMLs has not been clearly elucidated so far.

To characterize the methylation landscape of *DNMT3A*^INS^ AML in terms of methylation levels, we first examined whether *DNMT3A*^INS^ AMLs also show the hypomethylation patterns observed in *DNMT3A*^R882^ AMLs using Li2016 cohort, thereby seeking the similarities and differences of *DNMT3A*^INS^ and *DNMT3A*^R882^ methylomes. To determine the genomic regions subjected to *DNMT3A*^R882^-associated hypomethylation, we identified differentially methylated regions (DMRs) between *DNMT3A*^R882^ and *DNMT3A*^WT^ samples using an established method^18^. As expected, the identified DMRs predominantly consisted of hypomethylated DMRs (hypo-DMRs) in *DNMT3A*^R882^, accounting for 88% (465 of 527) of them (Figure 2a). Strikingly, we observed *DNMT3A*^INS^ AMLs showed comparable DNA methylation level to that of *DNMT3A*^WT^ at those identified *DNMT3A*^R882^-associated hypo-DMRs (Figure 2b). Additionally, these significant differences between *DNMT3A*^INS^ and *DNMT3A*^R882^ were also observed in TCGA-LAML and SNUH cohort (Supplementary Figure 5a and b). These results show that methylomes of *DNMT3A*^INS^ AMLs are devoid of *DNMT3A*^R882^-associated hypomethylation patterns and underscore the clear difference between *DNMT3A*^INS^ and *DNMT3A*^R882^ in terms of their methylomes.

**Figure 2.**
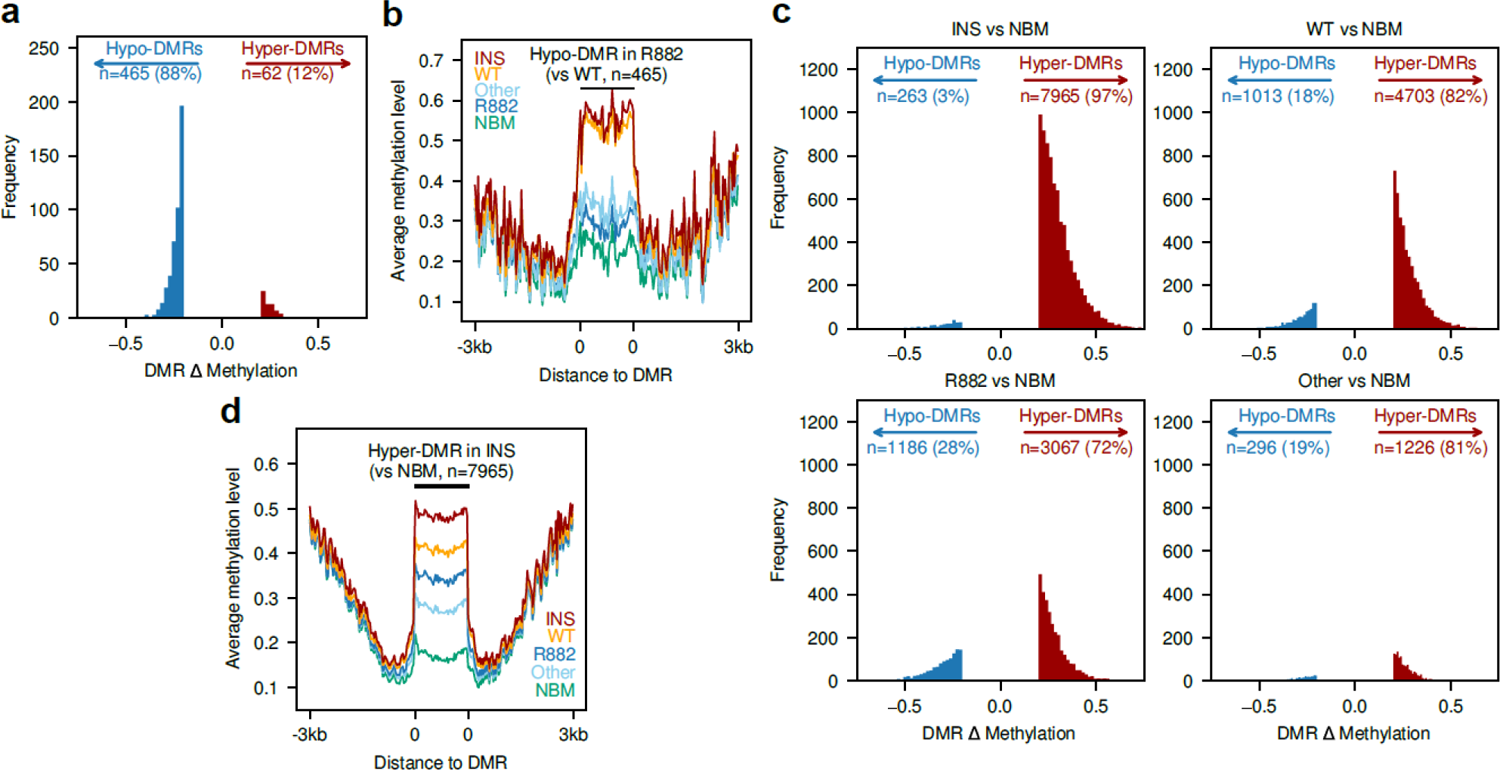
Methylomes of *DNMT3A*^INS^ AMLs are similar to *DNMT3A*^WT^ AMLs, but not to that of *DNMT3A*^R882^. (a) Distribution of average methylation level difference in DMRs identified between *DNMT3A*^R882^ and *DNMT3A*^WT^ AMLs. (b) Average methylation levels of different *DNMT3A* mutation subclasses of AMLs around the hypo-DMRs identified between *DNMT3A*^R882^ and *DNMT3A*^WT^ AMLs. (c) Distribution of average methylation level difference in DMRs identified between different DNMT3A mutation subclsses and normal bone marrow cells using RRBS. (d) Average methylation levels surrounding the hyper-DMRs in *DNMT3A*^INS^ (vs normal bone marrow cells) for each *DNMT3A* mutation subclass.

We were curious whether *DNMT3A*^INS^ AMLs harbor any regions having altered DNA methylation levels uniquely for them, so we identified and compared the characteristics of DMRs between each *DNMT3A* subclass and normal bone marrow (NBM) samples. As a result, *DNMT3A*^WT^ AMLs had 4703 (82%) hyper-DMRs and 1013 (18%) hypo-DMRs (Figure 2c). We note that the extreme bias toward hyper-DMRs may be due to a high specificity of eRRBS experiment for CpG-dense regions, which thus exaggerates cancer-associated hypermethylation events. Nevertheless, DMRs in *DNMT3A*^R882^ AMLs were less skewed toward hyper-DMRs. They were associated with fewer hyper-DMRs (n=3067, 72%) and more hypo-DMRs (n=1186, 28%; Figure 2c), recapitulating the attenuated hypermethylation in *DNMT3A*^R882^. DMRs identified in *DNMT3A*^INS^ AMLs were even more skewed toward hyper-DMRs (n=7965, 97%; Figure 2c). However, those hypermethylation events do not occur specifically in *DNMT3A*^INS^, as every *DNMT3A* subclasses of AMLs showed significant hypermethylation within the hyper-DMRs identified in *DNMT3A*^INS^ (Figure 2d) and even within the hyper-DMRs that were exclusive to *DNMT3A*^INS^ (Supplementary Figure 5c). The hyper-DMRs were also similarly distributed across genomic contexts (Supplementary Figure 5d). These observations indicate that the majority of hypermethylation in *DNMT3A*^INS^-associated hyper-DMRs originates from hypermethylation events that are generally observed in AML.

Altogether, these results suggest two conclusions for the methylation landscape of *DNMT3A*^INS^ AML. First, since *DNMT3A*^INS^ AMLs did not show *DNMT3A*^R882^-associated hypomethylation patterns, the current leukemogenic model for *DNMT3A*^R882^ may not directly apply to *DNMT3A*^INS^ AMLs. Next, the methylome of *DNMT3A*^INS^ showing comparable levels of DNA methylation to *DNMT3A*^WT^ implies that there are underlying molecular aberrations associated with *DNMT3A*^INS^ other than the absolute DNA methylation level changes. This underscores the importance of the increased intratumoral DNA methylation heterogeneity, including the local disorder of DNA methylation, in *DNMT3A*^INS^ AML.

### Local disorder of DNA methylation in *DNMT3A*^INS^ AML occurs predominantly at bivalent domains

Even though the precise molecular mechanism still remains obscure, previous experimental validation demonstrated that DNMT3A-dependent hypermethylation in AML cells occurs mostly at bivalent chromatin domains^17^. To provide additional line of evidence supporting that the observed hyper-DMRs in *DNMT3A*^INS^ truly resulted from the catalytic activity of DNMT3A, we took advantage of a reference epigenome of CD34+ myeloid progenitor from ENCODE^19^ and analyzed the epigenetic context of the hyper-DMRs. The resulting aggregated signals of several epigenomic marks surrounding the hyper-DMRs in *DNMT3A*^INS^ are shown in Figure 3a. We observed that these regions colocalized with both active (H3K4me1/3) and repressive (H3K27me3) histone marks, which indeed are indicative of bivalent chromatin domains. We additionally validated that the hyper-DMRs in *DNMT3A*^INS^ were strongly enriched for bivalent chromatin states inferred by ChromHMM^20^ (Figure 3b). Of note, the observed hypermethylation patterns enriched at bivalent domains are not restricted to *DNMT3A*^INS^, but also shown in all the other *DNMT3A* subclasses (Figure 3c, Supplementary Figure 6a), whereas hypo-DMRs were enriched for enhancer-related genomic contexts (Supplementary Figure 6b). Altogether, these data collectively indicate that the identified hyper-DMRs, primarily located at bivalent domains, represent the genomic regions where the *de novo* methylation by DNMT3A takes place.

**Figure 3.**
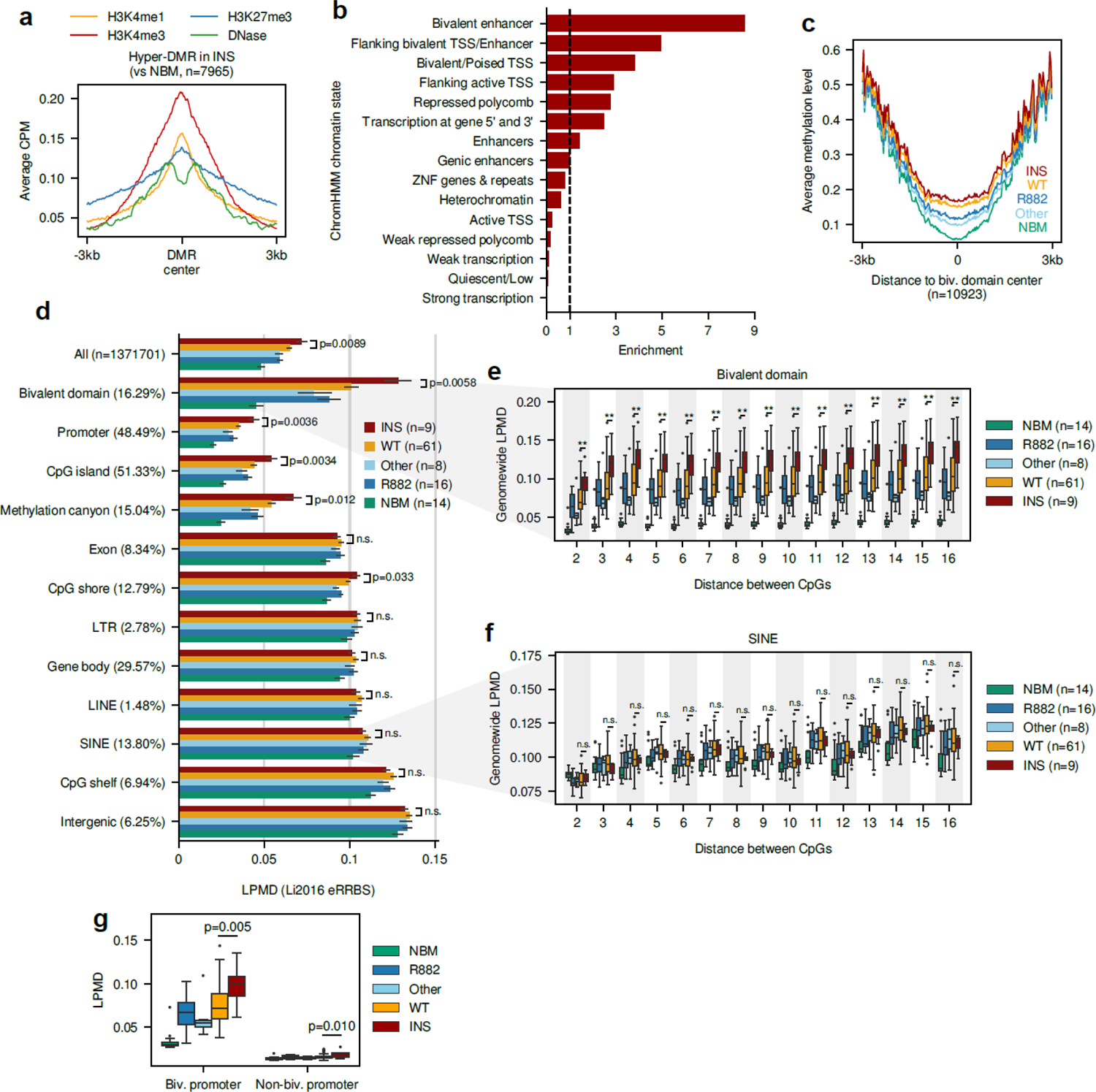
Local disorder of methylation in *DNMT3A*^INS^ AML occurs predominantly at bivalent domains. (a) Average histone modification levels around hyper-DMR identified between *DNMT3A*^INS^ and normal bone marrow cells. (b) ChromHMM chromatin context enrichment of hyper-DMR identified between *DNMT3A*^INS^ and normal bone marrow cells. (c) Average methylation level surrounding bivalent domains. (d) LPMD distribution across different genomic contexts for Li2016 cohort. Parenthesized values denote the proportion of the analyzed CpGs within each genomic context, except for that next to ‘All’, which denote the total number of analyzed CpGs. P-values from two-sided Mann-Whitney U tests between *DNMT3A*^WT^ and *DNMT3A*^INS^ are shown. Error bars denote standard errors. (e, f) LPMD distributions in (e) bivalent domain and (f) SINE. (g) LPMD comparison in bivalent or non-bivalent promoters. P-values from two-sided Mann-Whitney U tests between *DNMT3A*^WT^ and *DNMT3A*^INS^ are shown. In (e) and (f), ** p < 0.01, * p < 0.05, n.s. p > 0.05, two-sided Mann-Whitney U test. Throughout (e-g), the center line denotes the median, the upper and lower box limits denote upper and lower quartiles, and the whiskers denote 1.5× interquartile range.

Given that the bivalent domains are the putative hotspots of *de novo* methylation in *DNMT3A*^INS^ AMLs, we hypothesized that the DNA methylation disorder within those samples will be highly concentrated in those regions. To address this question, we computed LPMDs separately for 12 different genomic contexts. Surprisingly, we found that the difference of LPMD between *DNMT3A*^INS^ and the other DNMT3A subclasses was almost exclusive at bivalent domains and regulatory regions including promoters, CpG islands, shores, and methylation canyons (Figure 3d). This high specificity of DNA methylation disorder toward bivalent domain (Figure 3e) is notable when compared with the LPMD distributions for CpGs located at SINEs (Figure 3f). Note that those two genomic contexts harbor a comparable number of analyzed CpGs (223,428 and 189,338 CpGs for bivalent domains and SINEs, respectively). Further, categorizing promoters into bivalent and non-bivalent promoters revealed that the difference of LPMD was restricted to bivalent promoters, whereas non-bivalent promoters showed only marginal absolute difference of LPMD (Figure 3g). Taken together, we concluded that the disordered methylation in *DNMT3A*^INS^ AMLs is highly specific to bivalent domains, where the DNMT3A-driven *de novo* methylation potentially takes place. For convenience, we hereafter refer to the LPMD at bivalent domains as bivLPMD.

### DNA methylation disorder in *DNMT3A*^INS^ AMLs leads to increased epigenetic diversity of leukemic cell population

Our observations so far demonstrate that *DNMT3A*^INS^ AMLs were associated with the corruption of the local concordance of DNA methylation states. However, it should be interpreted with caution since it does not indicate the increase of the population-wise epigenetic diversity. LPMD is an intra-molecule measure^21^ that individually accounts for each read originated from a single cell, so it is not suitable to discern whether the erosion of local correlation of DNA methylation states occurs in a coordinated or stochastic manner throughout the malignant cells.

To determine whether the local disorder *DNMT3A*^INS^ AMLs accompanies the diversification of population-level epigenetic states, we orthogonally examined an inter-molecule DNA methylation heterogeneity score named epipolymorphism^22^. As a result, we observed significant increases of epipolymorphism in *DNMT3A*^INS^ AMLs (Figure 4a), indicating that the erosion of local concordance of DNA methylation in *DNMT3A*^INS^ AML occurs rather stochastically, and thus gives rise to the epigenetically diversified cell population. Of note, sample purity (Supplementary Figure 7a) and heterogeneity of cell type composition did not seem to confound the observed increased epigenetic diversity (Supplementary Figure 7b).

**Figure 4.**
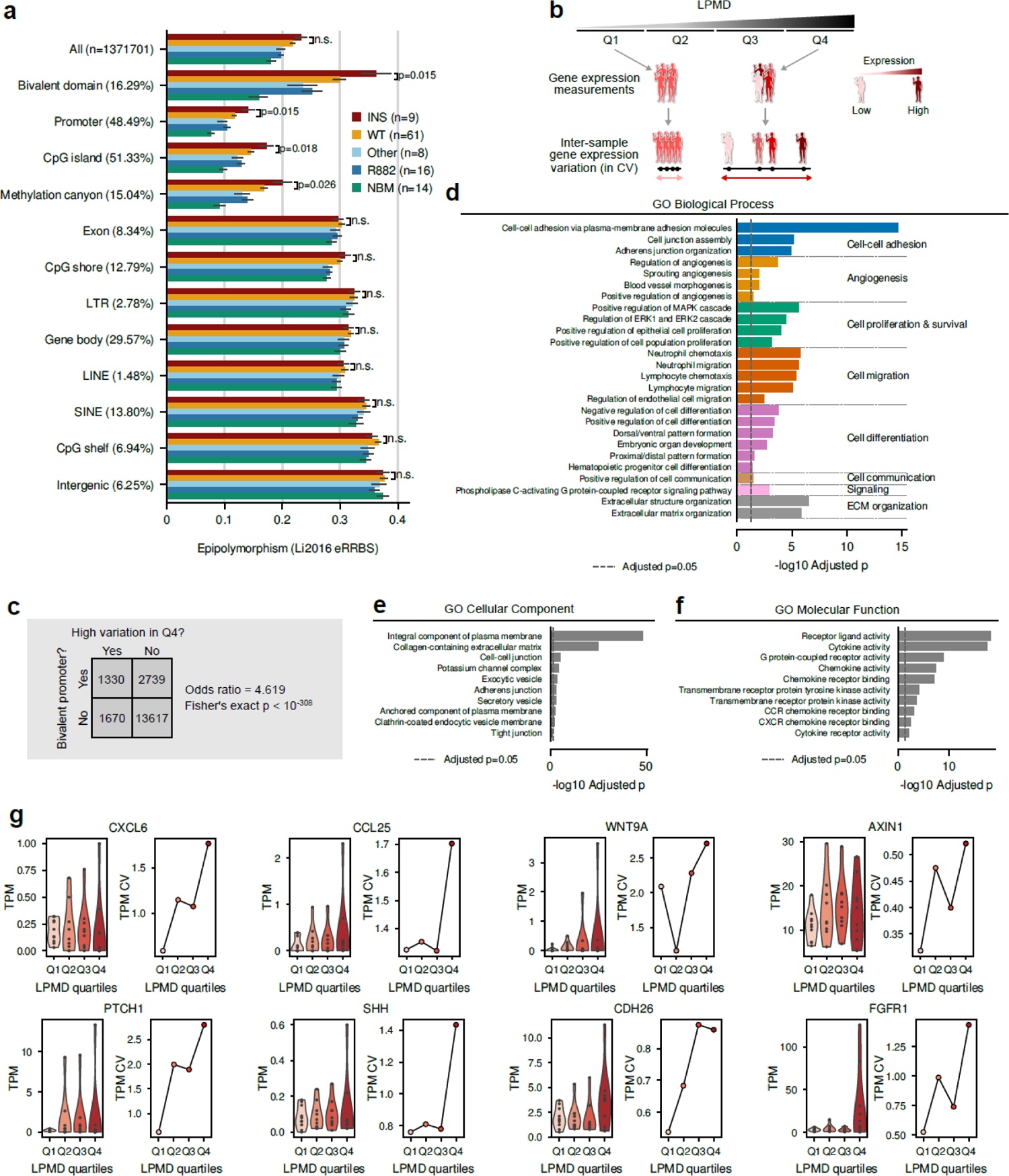
Functional implications of local disorder of DNA methylation and concomitant epigenetic diversity in AML. (a) Epipolymorphism distribution across different genomic contexts. P-values from two-sided Mann-Whitney U tests between *DNMT3A*^WT^ and *DNMT3A*^INS^ are shown. Error bars denote standard errors. (b) Experimental scheme to identify genes with methylation disorder-associated inter-sample expression variation. (c) Association between promoter bivalency and variable gene expression. Values in the table denote the number of genes in the corresponding condition. Odds ratio and p-value from two-sided Fisher’s exact test are shown. (d-f) Functional enrichment of top 4000 genes showing highest inter-sample expression variation in fourth quartile (Q4) of LPMD values for (d) GO Biological Process, (e) GO Cellular Component, and (f) GO Molecular Function terms. In (d), GO terms are grouped by broader biological concepts that are shown on the right side. (g) Gene expression levels (in TPM) and their inter-sample coefficient of variation (CV) for eight representative genes. Samples were grouped according to LPMD quartiles; Q1 (n=10), Q2 (n=9), Q3 (n=9) and Q4 (n=10).

### High LPMD is associated with increased transcriptional variance of genes involved in remodeling of leukemic stem cell niche

Given the remarkable specificity of local DNA methylation disorder and epigenetic diversity at bivalent domains, we then sought the functional implications of DNA methylation disorders in leukemia development at the transcriptome level. Importantly, the promoters of the developmental genes in stem cells are widely known to be frequently marked by bivalent chromatin marks^23^. Thus, the heterogeneity of DNA methylation in developmental promoters occurring at *DNMT3A*^INS^ AMLs suggests the possibility that the heterogeneity of the developmental gene regulation within leukemic cell population facilitates the progression of the disease by conferring the fitness advantage of cells.

To assess whether the epigenetic diversity of bivalent domains is associated with transcriptional diversity of the corresponding genes, a subset of Li2016 AML samples (n=38) profiled with both RRBS and RNA-seq data was analyzed. According to the additive property of variance, we assumed that the cell-level transcriptional variability, if it exists, will in turn manifest itself in the sample-level (i.e., bulk cell-level) transcriptional variability. Therefore, we measured and compared inter-sample variances of gene expression levels within each quartile of samples sorted by bivLPMD levels (Figure 4b).

We found that top 4,000 genes showing increased transcriptional variability in high-LPMD group (the highest quartile) were greatly enriched for genes having bivalent domains in their promoters (Odds ratio=4.619, p < 10^-308^, Fisher’s exact test; Figure 4c), which supports the linkage between the observed epigenetic heterogeneity of bivalent domains and the transcriptional heterogeneity. As expected, functions of those genes were enriched for cell differentiation (Figure 4d). Interestingly, we also found that they were also enriched for the biological processes shaping the hematopoietic stem cell niche in the bone marrow, including cell-cell adhesion, angiogenesis, cell proliferation and survival, cell communication, chemokine-mediated signaling and extracellular matrix organization (Figure 4d). Moreover, genes associated with high transcriptional variability were predominantly associated with cell membrane and extracellular matrix (ECM) (Figure 4e), suggesting the combinatorial diversification of the membrane protein configuration of progenitor cell, and eventually, the diversification of the modes of cell-cell and cell-ECM interaction within the hematopoietic stem cell niche. The enrichment of their molecular function towards membrane receptors, cytokines as well as chemokines also supports this notion (Figure 4f). Figure 4g demonstrates representative genes implying the heterogeneity of factors sculpting stem cell niche in high-bivLPMD AML samples. It highlights the transcriptional variability of cell adhesion molecule (*CDH26*), chemokines (*CXCL6* and *CCL25*), secreted signaling factors (*WNT9A* and *SHH*), signaling receptors (*PTCH1* and *FGFR1*) and downstream regulator (*AXIN1*). As *WNT9A* and *AXIN1* imply the heterogeneity of the activity of WNT signaling pathways, whose significance has been underscored in hematopoietic stem cell maintenance^24, 25^, we can envision that the diversity of the local concentration of paracrine factors in bone marrow stem cell niches may increase the fitness of leukemic stem cells communicating with it.

Collectively, these results showing the association of increased epigenetic and transcriptional variability propose a leukemogenic model that is worth exploring through functional experiments. It suggests that the increased transcriptional variability for both cell-intrinsic biological processes involving the balance between self-renewal and differentiation and cell-extrinsic factors surrounding each blast cell^26^ may confer fitness advantages to leukemic cells. Specifically, the external factors include direct interaction with other blast cells sharing the niche through cell-cell junctions, and other secretory factors including signaling molecules, cytokines and chemokines, produced by nearby cells triggering the intracellular signal transduction. A population of malignant cells experiencing locally heterogeneous environment may result in the increased adaptive potential of the disease.

### DNA methylation disorder at bivalent domains, but not absolute level of DNA methylation, robustly predicts the response of AML cells to hypomethylating agents

We then asked whether the local disorder of DNA methylation patterns at bivalent domains and associated epigenetic/transcriptomic diversity actually contribute to the sustained survival of AML cells. To examine the dependency of leukemic cells to DNA methylation disorder, we took a functional epigenomic approach by examining the survival of AML cells upon the elimination of the disorder of DNA methylation. To this end, we utilized the DNA methylation profiles of AML cell lines in Cancer Cell Line Encyclopedia (CCLE) and associated drug response profiles. Specifically, the drug responses of CCLE cell lines were collected from Cancer Therapeutics Response Portal (CTRP) v2, and DNA methylation profiles of corresponding cell lines were obtained by processing publicly available RRBS data by our own pipeline.

Meanwhile, hypomethylating agents (HMAs) including decitabine and azacitidine have been an invaluable epigenetic treatment option for AML patients who are not suitable for intensive chemotherapy^27^. Recent studies have shown complex and pleotropic mechanism of action of HMAs^28–30^, which in part explains why a robust biomarker predicting the response of a patient to HMA treatment still remains obscure. By examining the correlation between DNA methylation disorder and response of AML cell lines to HMA, we aimed to show the importance of the sustained methylation disorder in the survival of AML cells, as well as the potential of DNA methylation disorder as a biomarker for the response to HMA.

Strikingly, we observed a significant negative correlation between bivLPMD and the area under dose-response curve (AUDRC) of AML cell lines measured for decitabine (Figure 5a and b). This association persisted even when sufficient concentration of decitabine was treated in combination with other drugs (Figure 5a), suggesting that higher degree of DNA methylation disorder at bivalent domains predicts better response to decitabine. We additionally found that a high bivLPMD is also a good predictor of the response to RG-108, a non-nucleoside DNMT inhibitor that induces demethylation through direct binding to the active site of DNMTs (Figure 5a and b). We note that we could not observe any notable response to azacitidine for these AML cell lines, which may be due to an experimental artifact (Supplementary Figure 8). The association gradually diminished when the genomic regions for which LPMD values were calculated became distant from the core regulatory regions (from promoters and CpG islands to CpG shelves; Supplementary Figure 9), implying that the functional importance of the DNA methylation heterogeneity for the survival of AML cells was mediated by gene regulation. Remarkably, LPMDs calculated for non-bivalent non-regulatory regions did not show significant correlation with responses to HMAs (Figure 5c and d, Supplementary Figure 10) which further highlights that the DNA methylation disorder at bivalent regulatory domains is specifically important for the survival of AML cells. We also confirmed that bivLPMD did not correlate with the age of cell line at sampling time (Supplementary Figure 11a).

**Figure 5.**
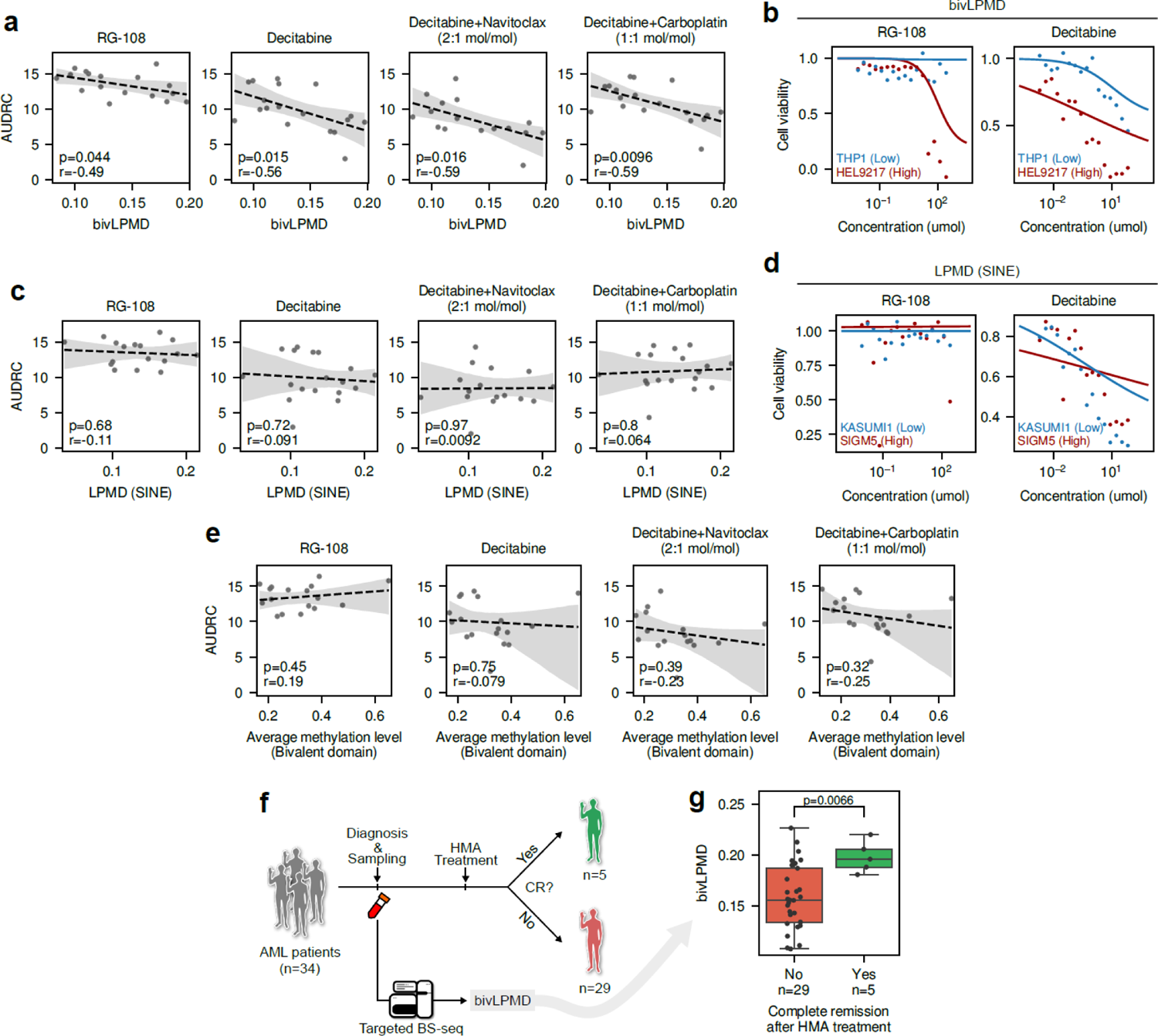
DNA methylation disorder at bivalent domain correlates with HMA responses of AML cells. (a) Correlation between LPMD at bivalent domains (bivLPMD) and area under dose-response curve (AUDRC) for hypomethylating agents. Pearsons’s correlation coefficients and corresponding p-values are shown. (b) Example dose-response curves for RG-108 and decitabine for two representative cell lines, THP1 and HEL9217, with low bivLPMD and high bivLPMD, respectively. (c) Correlation between LPMD at SINE and AUDRC for hypomethylating agents. Pearsons’s correlation coefficients and corresponding p-values are shown. (d) Example dose-response curves for RG-108 and decitabine for two representative cell lines, KASUMI1 and SIGM5, with low and high LPMD at SINE, respectively. (e) Correlation between average methylation level at bivalent domain and AUDRC. Pearsons’s correlation coefficients and corresponding p-values are shown. (f) Schematic diagram showing the retrospective analysis examining the utility of bivLPMD as a biomarker predicting hypomethylating agent (HMA) response. (g) Comparison of bivLPMD values in AML patient groups showing complete remission or not after HMA treatment. The center line denotes the median, the upper and lower box limits denote upper and lower quartiles, and the whiskers denote 1.5× interquartile range. P-value from two-sided Mann-Whitney U test is shown. BS-seq, bisulfite-sequencing.

Importantly, the responses of AML cell lines to decitabine and RG-108 were not associated with their methylation levels *per se* (Figure 5e). These results provide additional evidence supporting that focal increase of average methylation levels observed in AML is a mere collateral consequence of myeloproliferation, and the viability of AML cells generally does not depend on them. It is noteworthy that these results collectively suggest that AML cells were ‘addicted’ to the methylation disorder, since the erasure of disordered methylation states with hypomethylating agents triggered their death. To additionally confirm that our results on AML cell lines can be extended to clinical applications, we retrospectively measured the bivLPMD values using targeted enzymatic methyl-seq (EM-seq) from blood samples of 34 AML patients (Supplementary Table 3) who later underwent HMA treatment and examined its association with the response to HMA treatment (Figure 5f). Custom sequencing panel covering bivalent domains was designed for efficient measurement of bivLPMD through targeted EM-seq (Methods). Reassuringly, bivLPMD values were shown to be a good predictor of complete remission after HMA response (p=0.0066, two-sided Mann-Whitney U test; Figure 5g), while being not correlated with patient age (Supplementary Figure 11b). Collectively, these results show the importance of bivLPMD in the survival of AML cells, which is presumably due to the increased fitness advantage.

### Clinical implications of *DNMT3A*^INS^ in hematological disorders

Given the association between *DNMT3A*^INS^ and increased local methylation disorder and its functional impact in AML, we sought for the clinical outcomes of hematological conditions associated with *DNMT3A*^INS^. We first asked whether *DNMT3A*^INS^ is generally associated with adverse outcome of AML patients. To this end, we performed a pooled survival analysis of 668 non-M3 AML patients using three large cohorts (Ley et al. (n=233), TCGA-LAML (n=179) ^12^ and BeatAML (n=256) ^31^). Both *DNMT3A*^INS^ and *DNMT3A*^R882^ showed significantly poorer overall survival compared to *DNMT3A*^WT^ (log-rank p=0.0094 and 0.0047, respectively; Figure 6a), while *DNMT3A*^Other^ did not (p=0.482). Additionally, multivariate Cox regression showed that *DNMT3A*^INS^ is an independent risk factor (Hazard ratio 1.85, 95% CI 1.28-2.67) of AML even after accounting for age and gender (Figure 6b).

**Figure 6.**
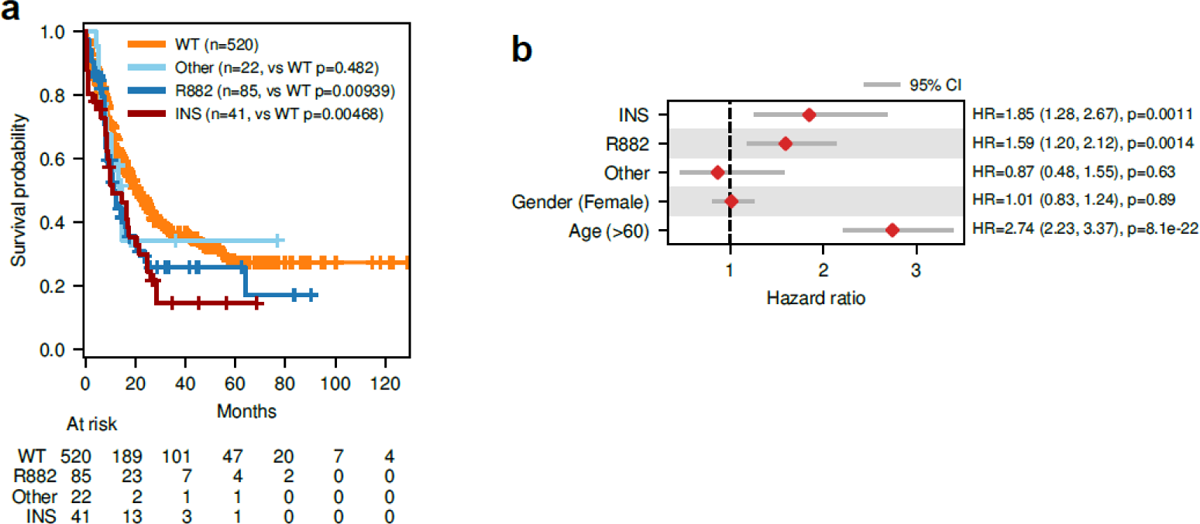
Clinical implication of *DNMT3A*^INS^ in hematological conditions. (a) Survival analysis for AML patients with different *DNMT3A* mutation subclasses. (b) Multivariate Cox proportional-hazards regression results.

## Discussion

AML comprises heterogeneous subtypes of diseases that can be classified under microscopic inspection of cells or based on genetic abnormalities. Although such subclassifications have been routinely utilized for the stratification of patient outcomes and the decision of treatment regimens, there is still enough room for the discovery and definition of further substratification of the disease. Since the early studies, the molecular classification of AML has highlighted remarkable recurrence of mutations in epigenetic modifiers including *DNMT3A*, *IDH1/2*, and *TET2*. However, the link between epigenetic alterations and aberrant epigenetic profiles has been only recently studied for its clinical relevance^11, 32^. In this regard, the complicated mutational landscape of *DNMT3A* involving conspicuous enrichment of mutations at residue R882 and dispersed mutations throughout non-R882 residues provides an excellent opportunity to investigate the mechanistic connection between genetic and epigenetic alterations.

In this study, we characterize the methylomes of AMLs harboring DNMT3A mutations that reduce the stability of the protein by analyzing the methylation profiles from three different AML cohorts. We show that they were associated with highly disordered local DNA methylation patterns specifically at bivalent domains, which in turn leads to the epigenetic diversity of AML cell population. As far as our knowledge is concerned, this is the first study that systematically analyzes the effect of the destabilization of DNMT3A directly on the methylomes of AML patients.

To date, researchers have been struggling to clearly provide the common effect of non-R882 DNMT3A mutations on leukemia, as the functional consequences of non-R882 mutations vary widely for the activity of the mutant proteins^33^. In line with this challenge, our results suggest a new perspective: the effect of individual non-R882 mutation on enzymatic activity may not be critical, at least for *DNMT3A*^INS^ mutations. This is because a mutant DNMT3A harboring one of those mutations is prone to be degraded and thus would not actively participate in *de novo* methylation. Instead, our results suggest that the common consequence of *DNMT3A*^INS^ variants, namely the reduction of intracellular DNMT3A concentration, is a key factor affecting the initiation and progression of AML.

Nevertheless, it seems that some *DNMT3A*^INS^ variants, especially those residing in the tetramer interface, further strengthen the dimeric preference of the enzyme by hampering the tetramerization by weakening the interaction at the tetramerization interface. Our experimental results showing the predominant dimerization of R736S DNMT3A *in vitro* (Supplementary Figure 3) suggest that some non-R882 variants may further promote the dimeric preference of the enzyme. Such residues that can elicit the synergy between destabilization and interface effect include S714 (stability score 0.688), R729 (stability score 0.364), R736 (stability score 0.316), R749 (stability score 0.339), S770 (stability score 0.419) and R771 (stability score 0.527), and they are shown to be among the most frequently mutated residues in hematological malignancies following R882 (Supplementary Figure 12).

Our observations suggest a potential explanation for the enigmatic recurrence of *DNMT3A*^INS^ variants in AML that has been poorly accounted for. In particular, our results link the biochemical property of DNMT3A^INS^ and the local DNA methylation disorder in *DNMT3A*^INS^ AML (Figure 7). The reduced dosage of intracellular DNMT3A due to the instability-driven degradation of DNMT3A^INS^ may favor the dimerization of DNMT3A over its tetramerization, as supported by the experimental study showing that the DNMT3A oligomerization is determined by its concentration^15^. Thus, *DNMT3A*^INS^ AML may show prevalent dimer-driven distributive *de novo* DNA methylation, whereas *DNMT3A*^WT^ AML exerts tetramer-driven processive catalysis. Distributive methylation leads to a decreased concordance of local DNA methylation states, and the random dissociation of DNMT3A dimers from DNA in turn triggers the concomitant increase of the epigenetic diversity of cancer cell population. Although the clear mechanism of how the epigenetic diversity drives the progression and aggressiveness of AML cells still remains to be elucidated, our results showed the association between epigenetic and transcriptional heterogeneity of leukemic cells. Especially, the functional heterogeneity was enriched for genes contributing to the fitness of leukemic stem cells within the hematopoietic stem cell niche. Furthermore, the correlation between the epigenetic diversity at bivalent regulatory domains and response to HMA implies the connection between epigenetic diversity and transcriptional heterogeneity of cancer cells.

**Figure 7.**
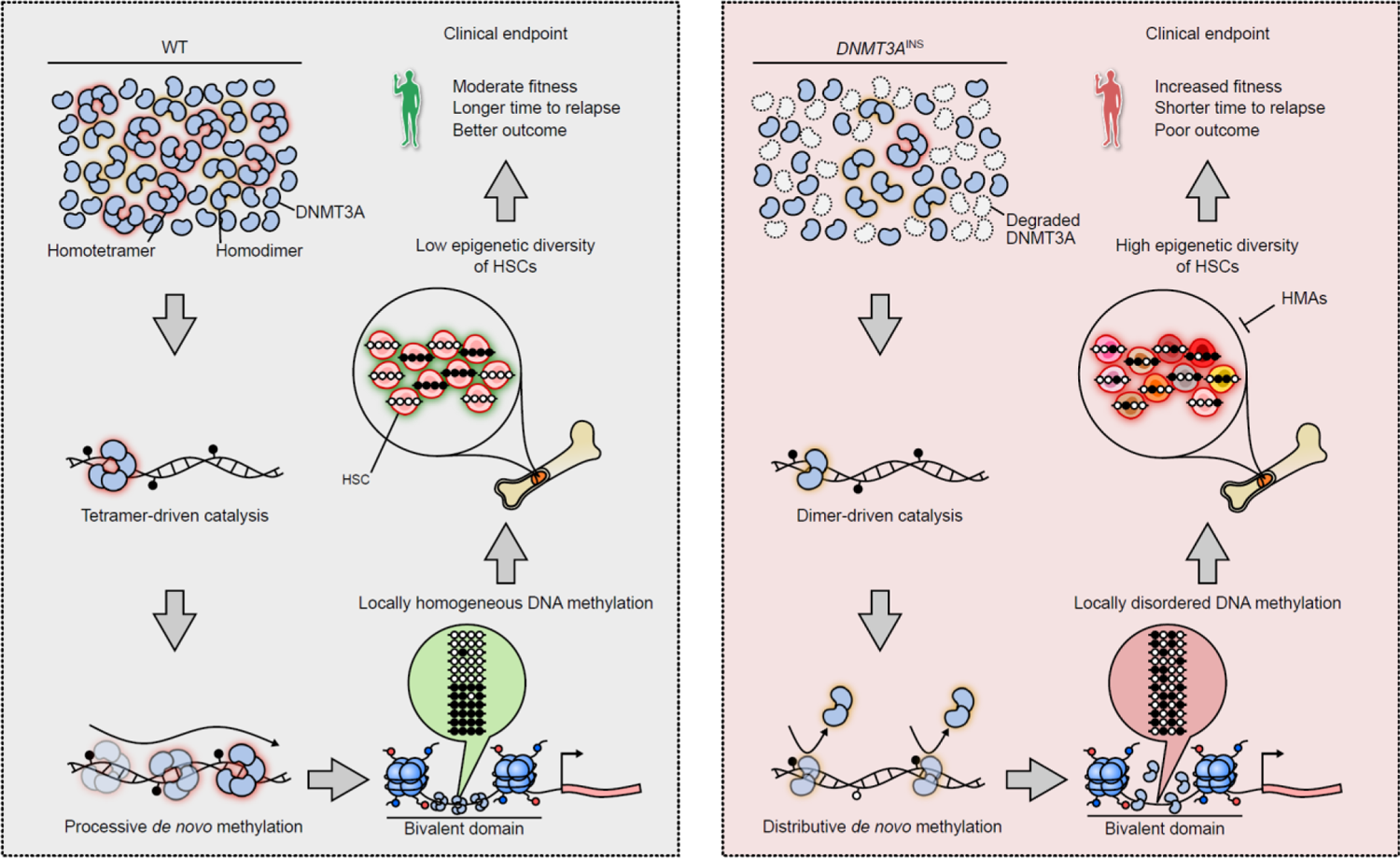
Proposed model explaining the DNA methylation disorder induced by mutant DNMT3A^INS^ and its clinical implications. Proteasomal degradation of destablized DNMT3A proteins harboring *DNMT3A*^INS^ mutations leads to decreased effective concentration of intracellular DNMT3A. Thus, the dimerization of DNMT3A protein is preferred over their tetramerization. Unlike DNMT3A tetramers, which conduct *de novo* methylation in a processive manner, DNMT3A dimers dissociate from DNA frequently during catalysis. This distributive *de novo* methylation results in stochastic local disorder of DNA methylation patterns, which in turn confers population-level epigenetic diversity of hematopoietic stem cells. Increased epigenetic diversity of cell population translates to increased fitness or adaptive potential of cell population, ultimately leading to poorer outcome of the patients.

Cancer has long been appreciated as an intrinsically heterogeneous disease. Genetically and epigenetically distinct cells, or subclones, arise from sporadic molecular aberrations, and they compete and cooperate with each other while exploiting the limited resources surrounding them. For recent decades, the extent of such intratumor heterogeneity has shown great potential as a clinical biomarker. However, studies so far have primarily focused on their prognostic power, and it is still questionable that the heterogeneity itself can be exploited as an actionable therapeutic target. In this regard, epigenetic intratumor heterogeneity, thanks to its reversible nature, would bring a novel therapeutic avenue that exploits direct manipulation of the heterogeneity of cancer cell population, i.e., homogenization of epigenetic states of cancer cells. Such intervention may undermine the fitness of cancer cell population, which ultimately triggers cell death. Indeed, this proposed mechanism may have already been implicitly functioning behind the conventional HMA treatments, but it has not been clearly elucidated before, as shown by the lack of DNA methylation-based biomarkers for HMAs. Our observations from functional epigenomic analyses in part support this scenario, and further provide an effective way to predict the response of AML cells to HMAs, which greatly increase the precision of the antileukemic therapies in clinical practice.

## Methods

### RRBS

To construct the MSP1 and Apek1 digested reduced-representation bisulfite sequencing (RRBS) library, 500 ng of input genomic DNA was assembled into 50 μl of reactions with MspI (NEB), incubated at 37°C for 24-26 h. ApeKI (NEB) was then added and incubated at 75°C for 16–20 h. The digested products were purified with a MiniElute PCR Purification Kit (Qiagen). After purification, the digested products were blunt-ended, and then dA was added, followed by methylated-adapter ligation. A range of 160-420 adapter-ligated fraction was excised from a 2% agarose gel. Bisulfite conversion was conducted using a ZYMO EZ DNA Methylation-Gold Kit™ (ZYMO), following the manufacturer’s instructions. The final libraries were generated by PCR amplification using PfuTurbo Cx Hotstart DNA polymerase (Agilent technologies, Santa Clara, CA, USA). RRBS libraries were analyzed by an Agilent 2100 Bioanalyzer (Agilent Technologies). The methylation data were generated using two different platforms, Illumina HiSeq 2500 Standard 100 PE (100bp paired end) and NovaSeq 6000 S4 150 PE (150bp paired end).

### Collecting and processing public DNA methylation data

DNA methylation profiles for the public cohorts analyzed in this study were collected and processed as follows. Raw eRRBS sequencing data for 47 AML patients^8^ were obtained from dbGaP under accession phs001027.v2.p1. Sequencing was performed for each patient at both points of diagnosis and relapse, thus resulting in 94 sequencing runs in total. Bisulfite sequencing reads were adapter-trimmed with Trim galore!^34^ v0.6.7 with --rrbs option turned on. Reads were aligned to the hg38 reference genome with Bismark^35^ v0.22.3, and CpG methylation levels were extracted using MethylDackel^36^ v0.4.0. The same RRBS processing pipeline was applied to our own SNUH cohort. Illumina HumanMethylation450 BeadChip array-based DNA methylation profiles of 140 TCGA-LAML patients were downloaded from Genomic Data Commons (GDC) data portal.

### Sample collection for SNUH cohort

The samples were collected in accordance with the guidelines and regulations of the Seoul National University Hospital [IRB No. H-1103-004-353]. DNMT3A mutations for patients with AML or myelodysplastic syndrome patients were identified using clinical NGS panel screening.

### Definition of *DNMT3A*^INS^ variants and *DNMT3A*^INS^ AML

*DNMT3A*^INS^ variants were identified using the catalog of stability ratios of *DNMT3A* amino acid substitutions that were experimentally determined by previous study^6^. Although the catalog covers a large number of residues (248 / 912 amino acids), still some of mutations occurring in clinical AML samples are not covered. Therefore, we extrapolated the ratios to assign stability scores for those uncharted substitutions by assigning a single stability score for each amino acid position, instead of each amino acid substitutions. It was done by computing the average of all known stability ratios resulting from the substitution each amino acid. Indeed, this procedure makes individual stability score less sensitive to the amino acid properties, thus some false positive or negative *DNMT3A*^INS^ classification can be produced. However, we considered that it will be more beneficial to increase the sensitivity of the whole study by incorporating more variants to the analyses.

All variants having processed stability scores below 0.75 were classified as *DNMT3A*^INS^. Moreover, nonsense and frameshift variants were also included as part of *DNMT3A*^INS^ variants, as the truncation of DNMT3A protein are known to cause protein degradation in AML cells^37^. An AML sample was classified as *DNMT3A*^INS^ AML only if it harbors a single mutation on *DNMT3A* gene and it is *DNMT3A*^INS^. If a sample harbor *DNMT3A*^R882^ mutation, it was classified as *DNMT3A*^R882^ AML regardless of the existence of other mutations to reflect the dominant-negative effect of *DNMT3A*^R882^ variant. All the other samples having non-destabilizing variants or multiple variants were classified as *DNMT3A*^Other^.

### Collecting and processing somatic mutation profiles

Somatic variants for each individual were determined as follows. Whole exome sequencing data for Li2016 cohort were accessed via dbGaP under accession phs001027.v2.p1. In total, whole exome sequencing runs for 94 cancer samples (diagnosis and relapse) as well as 47 matched normal samples were obtained. Reads were aligned to hg38 reference genome with bwa v0.7.17-r1188^38^. To increase the sensitivity of variant calls, we considered somatic variants called by at least one of Strelka2^39^ v2.9.10 and Varscan^40^ v2.4.4 as valid somatic variants. Resulting variants were annotated with SnpEff^41^ v5.0 and SnpSift^42^ v4.3t. Finally, variants were post-filtered to avoid false positive calls using the following criteria: (1) variants should be present with variant allele frequency greater than 5%, (2) variant alleles should be supported by at least five sequencing reads, (3) variants should not be present with ExAC population allele frequency more than 1%, and (4) only missense, nonsense, frameshift and splice variants were considered. For TCGA-LAML cohort, we collected the corresponding mutational profiles from cBioPortal^43^.

### Computation of local pairwise methylation discordance (LPMD)

To measure the disorder of DNA methylation, we devised a new measure called local pairwise methylation discordance (LPMD) that measures the extent to which a pair of nearby CpGs at a fixed distance have conflict in their methylation states. LPMD takes advantage of the phased methylation states of nearby CpGs that are simultaneously captured by a single bisulfite sequencing read. Through the enumeration of all the sequencing reads, LPMDd is computed as the proportion of CpG pairs at genomic distance *d* (in bp) with different methylation states. LPMD values were computed using Metheor v0.1.2^16^.

On the other hand, we cannot extract a pair of DNA methylation states that originates from a single cell (i.e., phased methylation states) using the results from DNA methylation arrays. To approximate sequencing-based LPMD values using methylation levels measured by DNA methylation arrays, the difference of DNA methylation levels of a CpG pair at a fixed distance was utilized. The use of this measure can be justified by the fact that the methylation level difference of CpG pair forms the lower bound of LPMD. Assume that there is a CpG pair with methylation level β_1_ and β_2_, where β_1_ < β_2_, without loss of generality. Then, the maximum proportion of CpG pairs both having methylated state will be β_1_. Similarly, the maximum proportion of CpG pairs both having unmethylated state will be 1 − β_2_. Thus, the lowest possible proportion of CpG pairs having different methylation state is 1 − (β_1_) − (1 − β_2_) = β_2_ − β_1_, which is the methylation level difference of the pair. Sample-wise array-based LPMD was computed similarly to sequencing-based LPMD by specifying the distance between CpG pairs.

### Computation of epipolymorphism

Epipolymorphism^22^ is a cell population-wise measure that quantifies the diversity of methylation patterns, or epialleles, of four consecutive CpG sites (CpG quartets). To compute epipolymorphism from bisulfite read alignments of Li2016 cohort, we only considered CpG quartets that are supported by more than ten sequencing reads. CpG quartets harboring CpG site that overlaps with dbSNP 151 SNPs were excluded. For each CpG quartets, epipolymorphism is defined considering 16 possible patterns of DNA methylation states. For convenience, here we denote unmethylated and methylated states as ‘0’ and ‘1’, respectively. Then we can think of 16 possible DNA methylation patterns from *x*_0_ = 0000 (fully unmethylated pattern) to *x*_15_ = 1111 (fully methylated pattern), and epipolymorphism is defined as below.

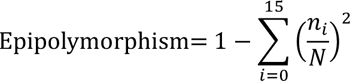

where *n*_*i*_ denotes the number of reads supporting pattern *x*_*i*_ and 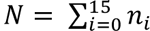. Epipolymorphism values were computed using Metheor v0.1.2^16^.

### Reference epigenome for CD34 hematopoietic stem cells

Reference epigenomes for CD34-positive hematopoietic stem cells (HSCs) were downloaded from ENCODE under accession number ENCSR970ENS. In particular, the raw whole genome bisulfite sequencing data was downloaded under library accession ENCLB590SRF and was processed as described above. Processed signal p-values and called peaks for ChIP-seq targeting H3K4me1, H3K4me3, H3K9me3, H3K27me3, H3K27ac and H3K36me3 histone marks were downloaded under accession number ENCSR401CJA, ENCSR136QKZ, ENCSR957WQX, ENCSR355PUX, ENCSR620AZM and ENCSR164ROX, respectively. Similarly, signal p-values and peaks for DNase I hypersensitive sites were downloaded under accession ENCSR468ZXN. For the subsequent analyses, signal p-values were normalized with arcsinh transformation. The core 15-state chromatin states inferred by ChromHMM^20^ were downloaded from Roadmap Epigenomics for the enrichment analysis of differentially methylated regions. Bivalent domains in CD34-positive hematopoietic stem cells are defined as the genomic regions with chromatin states named 10_TssBiv, 11_BivFlnk or 12_EnhBiv.

### Selection of the bivalent domains for targeted enzymatic methyl-seq

We selected representative bivalent domains that show pronounced methylation disorder in *DNMT3A*^INS^ AMLs compared to HSCs for targeted enzymatic methyl-seq (EM-seq). To obtain sufficient depths for the targeted regions, the total span of the sequencing panel was aimed to be about 500kbp, which is about 4% of the bivalent domains in the HSC reference epigenome (∼12,526 kbp in total). The following describes how we prioritized bivalent domains to be selected for the panel. First, bivalent domains were ranked by average difference of DNA methylation level between SNUH5763 sample and HSC reference epigenome. At the same time, they were ranked also by density of containing CpGs (number of CpGs divided by the length of the region). Of note, we found that a majority of (90%) bivalent domains were hypermethylated, and higher density of CpGs was positively correlated with methylation level difference (Pearson’s r=0.554, p < 10^-308^). Final ranks were obtained by taking geometric mean of methylation level difference and CpG density for each bivalent domain and the top 454 bivalent domains spanning 499,859bp were selected for the panel.

### Targeted enzymatic methyl-seq

We applied an improved methylation detection using EM-Seq to avoid loss of DNA, GC biased coverage, and poor complexity compared with BS-Seq^44^. Targeted capture panel was designed to tile the selected bivalent domains (10bp flanking). 4,200 hybrid capture probes using the Twist target enrichment (Twist Bioscience, San Francisco, CA, USA) were synthesized to capture ∼58,000 CpGs within the selected bivalent domain regions. Genomic DNA samples were fragmented physically by Covaris (200 to 300bp). Methylated cytosine residues of initial 200ng input gDNA were converted enzymatically by Twist Bioscience’s NEBNext Enzymatic Methyl-Seq (EM-Seq). Then pre-PCR amplification and sample library preparation were processed. Twist fast hybridization target enrichement with 8-plexing, post PCR amplification, and libraries were sequenced on DNBSEQ-G400 Dx (MGI Tech, Shenzhen, CHINA) with 100bp paired-end reads with a minimum coverage of 280x (average coverage 380x; 234x∼559x).

### Genome annotations

All the bioinformatics analyses were performed with hg38 human reference genome. Annotations for human CpG islands were downloaded from UCSC Table Browser. Based on the CpG island annotations, annotations for CpG shores (defined as up/downstream 2kb regions flanking CpG islands) and CpG shelves (defined as further up/downstream 2kb regions flanking the borders of CpG shores) were obtained using BEDTools^45^ v2.26.0. Gene annotations were obtained from GENCODE^46^ v32 release. Annotations for CpG methylation canyons were obtained from a previous study^47^.

### Identification of differentially methylated regions

Differentially methylated regions (DMRs) between various *DNMT3A* subclasses were identified by metilene v0.2-8^18^. We required at least 4 CpGs for a region to be called as a DMR, while allowing at most 500bp-away adjacent CpG pair within a DMR. Among those candidate regions, regions showing methylation difference greater than 0.2 and showing Benjamini-Hochberg adjusted p-value less than 0.01 were finally called as DMRs.

### Drug response analysis

Drug response analyses were conducted by reanalyzing public experimental results for Cancer Cell Line Encyclopedia (CCLE)^48^ cell lines. Only the cell lines of hematopoietic lineage derived from AML that have associated raw RRBS data were used. Raw RRBS data were obtained under SRA accession SRP186687 and processed as described above. To avoid spurious methylation calls we excluded CpGs that overlaps with SNPs using dbSNP version 151. Moreover, we excluded CpGs located at ENCODE blacklisted regions^49^ and their flanking 1000bp regions from analysis.

The responses of the cell lines to hypomethylating agents were adopted from Cancer Therapeutics Response Portal (CTRP) v2^50^. Area under drug response curve (AUDRC) was used as a measure of drug response, and the fitted curve was reconstructed and visualized with the following four-parameter logistic nonlinear regression model^51^:

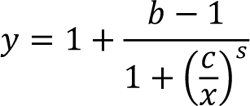

where *x* is the concentration of the drug at which the response of cells is to be computed, *c* is the dosage of the drug where the 50% of cells shows response, *b* is the baseline response, which denotes the response of cells at sufficiently high concentration of the drug and *s* is the steepest slope of the logistic curve.

## Acknowledgements

This research was supported by a grant from the Korea Health Technology R&D Project through the Korea Health Industry Development Institute (KHIDI), funded by the Ministry of Health & Welfare, Republic of Korea (grant number: HI18C1876) (to Y. K). This research was also supported by a grant (grant number: NRF-2020R1A2B5B03001517) from the National Research Foundation of Korea (to J.S). Genome Opinion Inc. implemented and optimized the targeted enzymatic methyl-seq panel (LifeEx EM) and provided the sequencing data.

## Additional information

All RRBS and EM-seq data generated in this study were deposited in the NCBI Sequence Read Archive (SRA) database under project accession PRJNA933381.

## Supplementary Figures

**Supplementary Figure 1.**
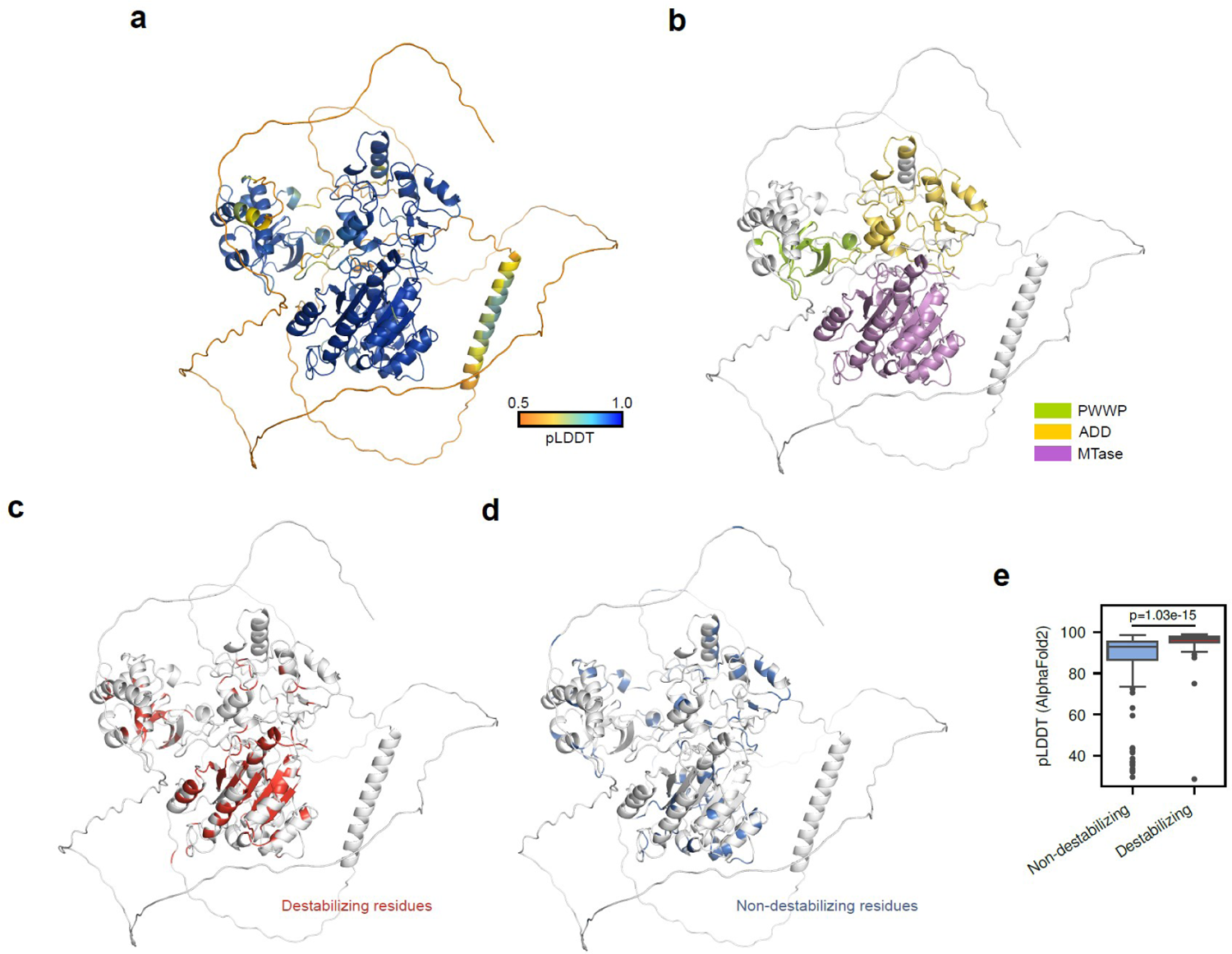
Structural properties of DNMT3A amino acid residues inducing protein instability upon mutation. Predicted structure of full-length DNMT3A was obtained from AlphaFold Protein Structure Database under Uniprot accession Q9Y6K1. (a) Residues were colored according to predicted local distance difference test (pLDDT) values produced from AlphaFold2 model. (b) Residues were colored according to the conserved domains. (c) Destabilizing residues (n=125) were colored in red. (d) Non-destabilizing residues (n=123) were colored in blue. (e) Boxplot showing the difference of pLDDT values between non-destabilizing and destabilizing residues. The center line denotes the median, the upper and lower box limits denote upper and lower quartiles, and the whiskers denote 1.5× interquartile range.

**Supplementary Figure 2.**
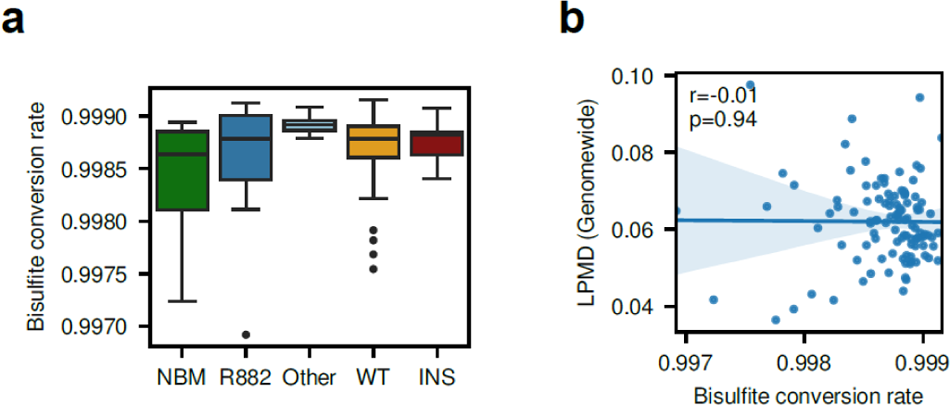
Bisulfite conversion rate did not affect the observed high LPMD in *DNMT3A*^INS^. (a) Boxplot showing the distribution of bisulfite conversion rate for each DNMT3A subclasses in Li2016 eRRBS data. (b) No correlation was observed between bisulfite conversion rate and genomewide LPMD values.

**Supplementary Figure 3.**
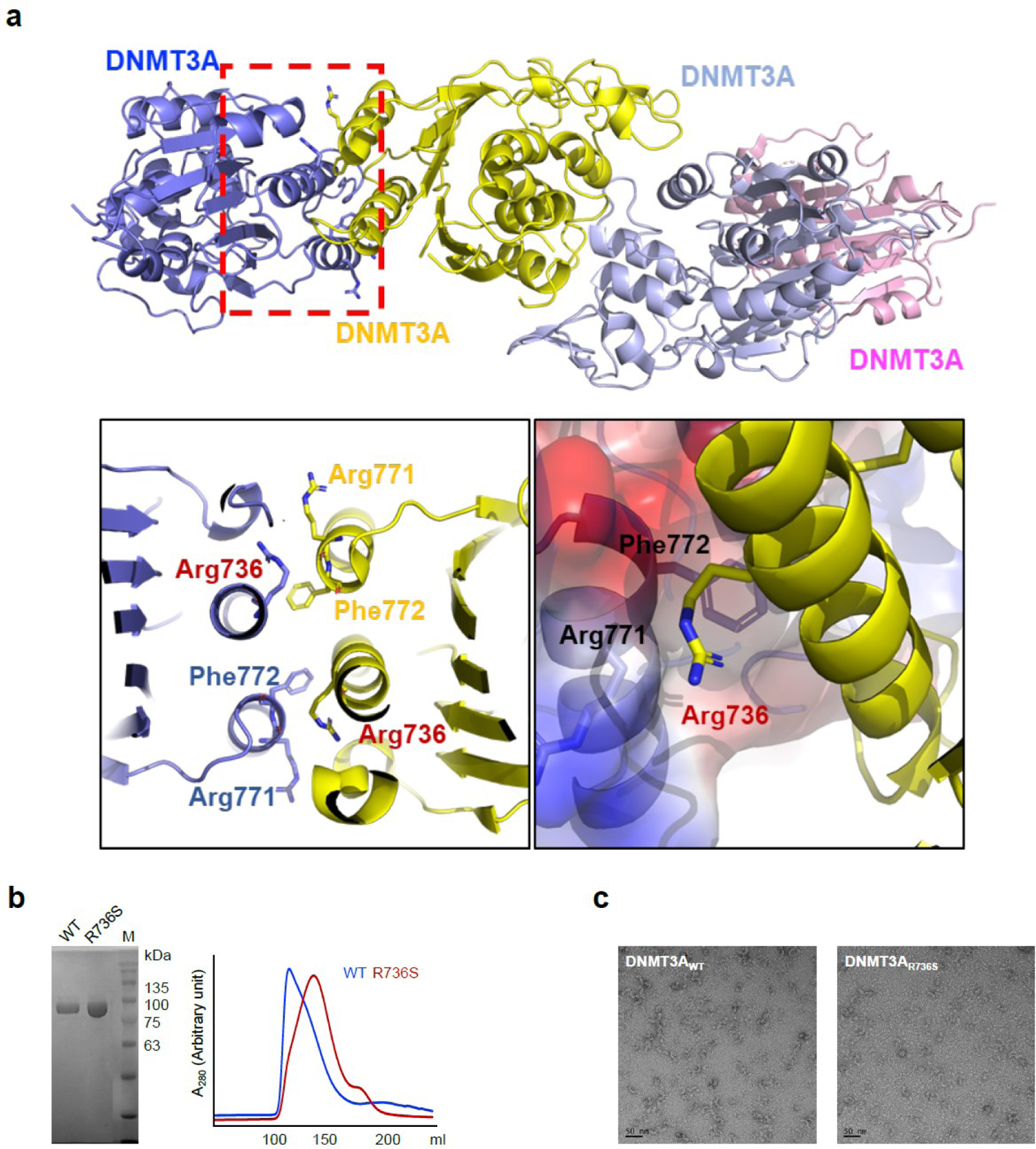
R736S mutation disrupts the oligomerization of DNMT3A. (a) DNMT3A tetramer status was modeled based on DMNT3A-DMNT3L heterotetramer (PDB ID 6BRR) by superimposing DMNT3A on DMNT3L. The location of R736S at the DNMT3A oligomer interface is indicated with a red dotted box. The detailed view of the interaction between Arg736 (R736) and the Arg771 and Phe772 from the adjacent DNMT3A molecule (left panel). The hydrocarbon region in the Arg771 side chain and the phenyl ring in Phe772 form a hydrophobic patch where the hydrocarbon region of Arg736 interacts. The adjacent DNMT3A molecule is shown in electrostatic surface representation, showing the hydrophobic interaction among Arg736, Arg771 and Phe772. The mutation of Arg736 would interfere this interaction. (b) An SDS-PAGE gel shows the purified DNMT3A^WT^ and DNMT3A^R736S^. A gel-filtration chromatograms of DNMT3A^WT^ (WT) and DNMT3A^R736S^ (R736S) shows that R736S disrupts the oligomerization states. (c) Negative-stain EM analysis of DNMT3A^WT^ and DNMT3A^R736S^. DNMT3A^R736S^ exhibits smaller particles than DNMT3A^WT^.

**Supplementary Figure 4.**
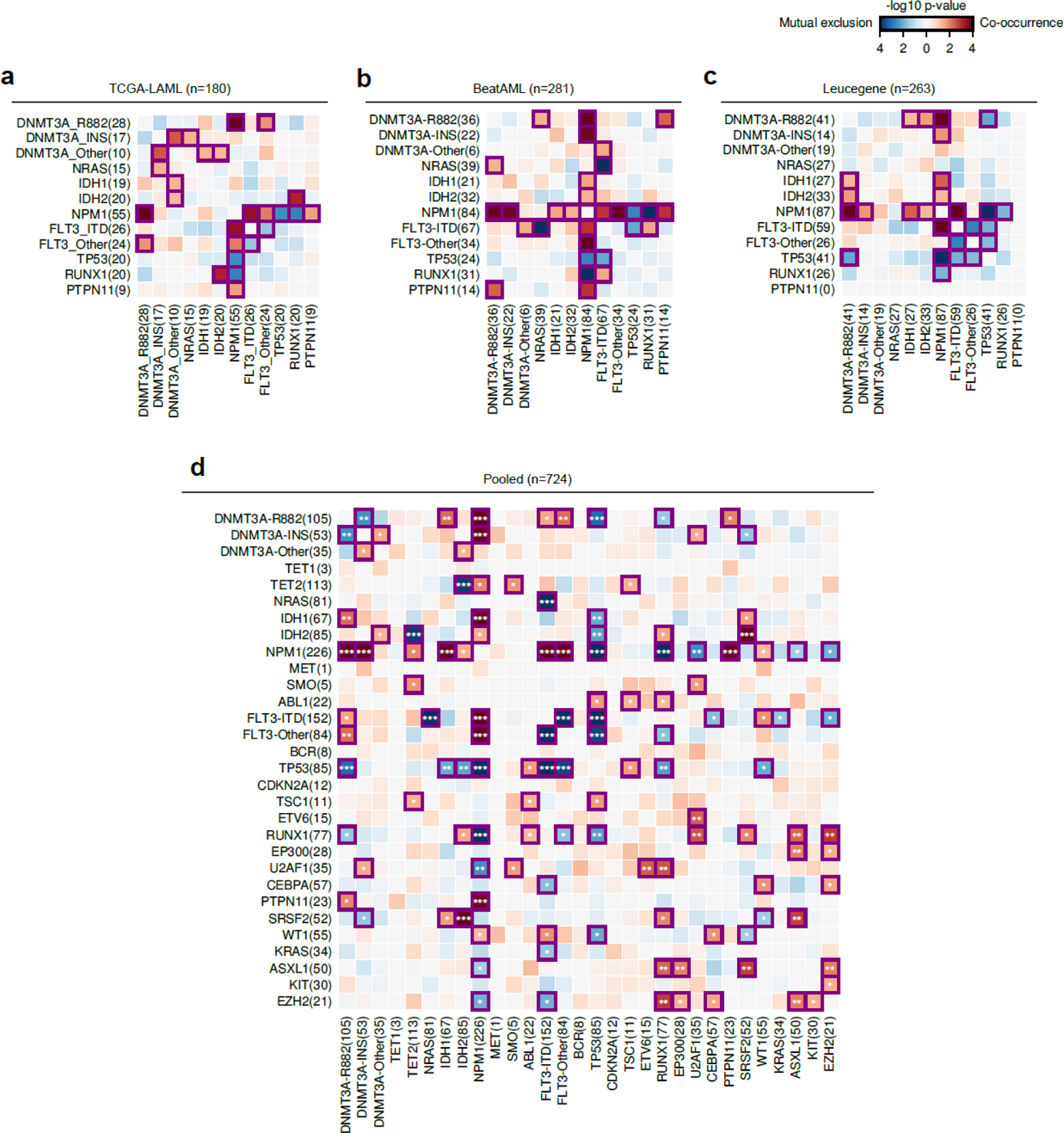
Mutational co-occurrence analysis. Heatmaps show the significance of the co-occurrence (red) and mutual exclusion (blue) of a pair of mutations for (a) TCGA-LAML, (b) BeatAML and (c) Leucegene cohorts. Colors denote unadjusted p-values from Fisher’s exact test. Gene pairs with p < 0.05 were indicated with purple squares. FLT3-ITD, FLT3 with internal tandem duplication; FLT3-Other, FLT3 with mutations other than FLT3-ITD.

**Supplementary Figure 5.**
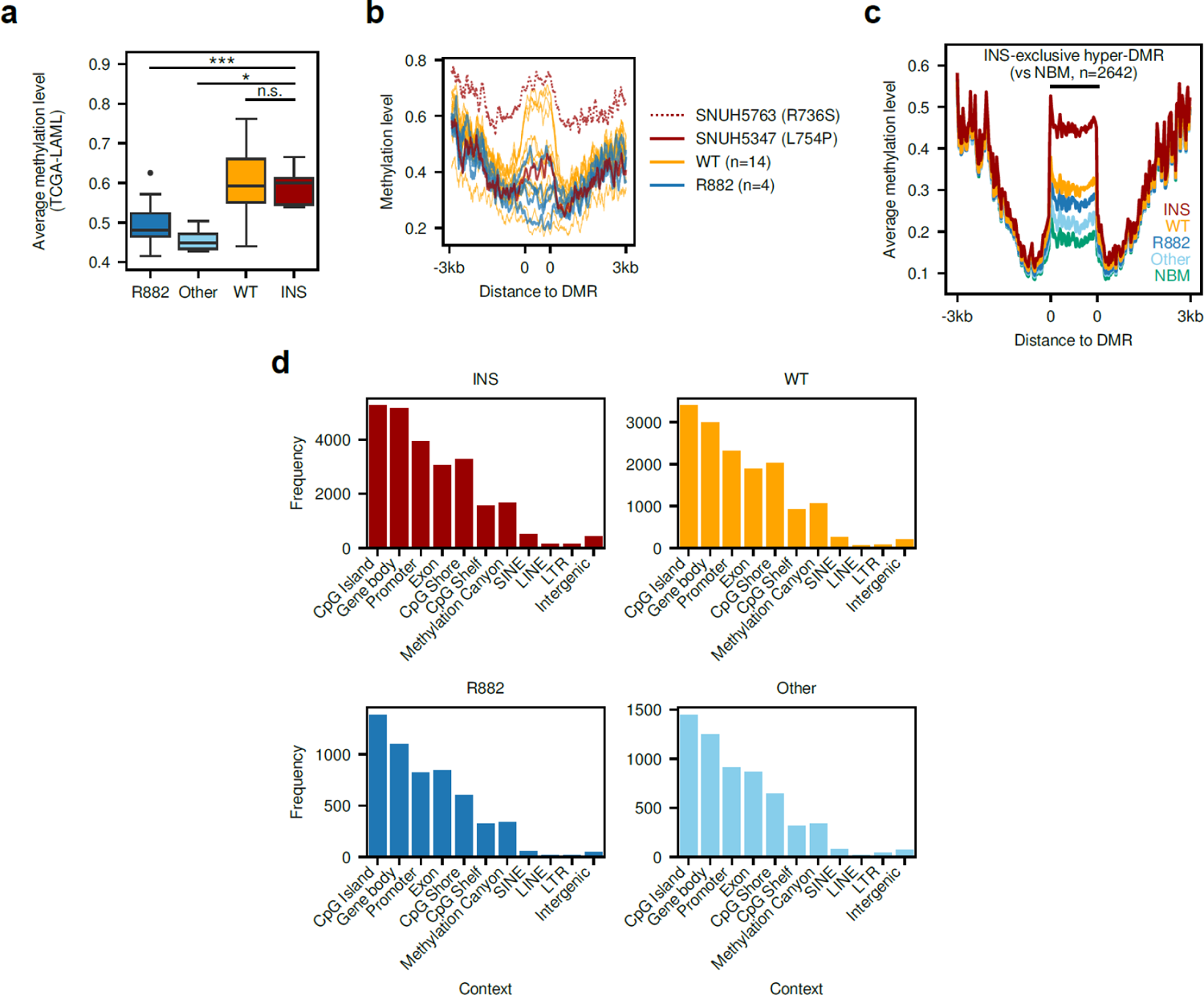
Differentially methylated region (DMR) analysis. (a) Average methylation levels of TCGA-LAML cohort samples within hypo-DMRs in *DNMT3A*^R882^ (vs *DNMT3A*^WT^) defined in Li2016 cohort. (b) Methylation levels of SNUH cohort samples surrouding hypo-DMRs in *DNMT3A*^R882^ (vs *DNMT3A*^WT^) defined in Li2016 cohort. (c) Average methylation levels of Li2016 cohort samples surrounding hyper-DMRs in *DNMT3A*^INS^ (vs normal bone marrow cells). (d) Frequencies of genomic contexts covered by hyper-DMRs (vs normal bone marrow cells) for each DNMT3A subclasses in Li2016 cohort. NBM, normal bone marrow.

**Supplementary Figure 6.**
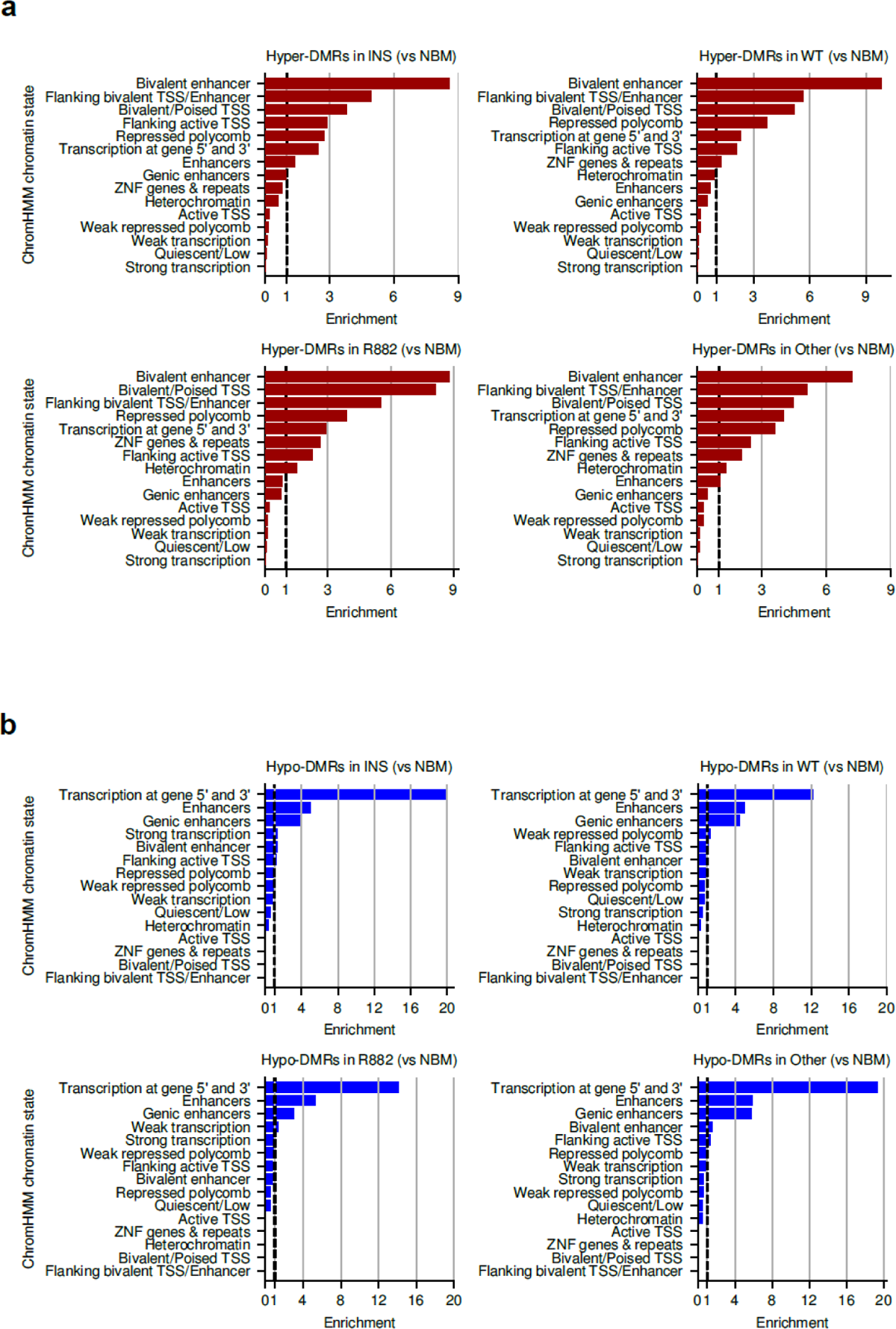
Testing enrichment of differentially methylated regions for chromatin contexts. (a) Fold enrichment of the occurrence of hyper-DMRs (vs normal bone marrow cells) for each chromHMM chromatin state. Fold enrichment was computed by taking the ratio between the length of the observed and expected intersection between DMRs and each chromatin state. (b) Fold enrichment of the occurrence of hypo-DMRs (vs normal bone marrow cells) for each chromHMM chromatin state.

**Supplementary Figure 7.**
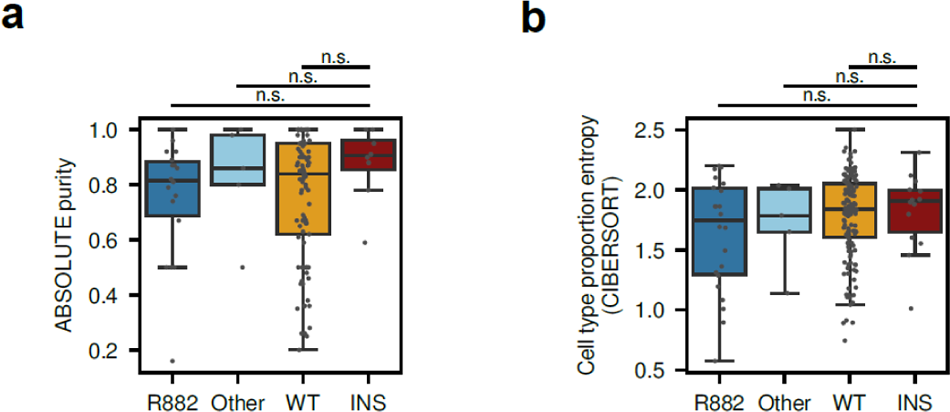
Purity and cell type composition analysis. (a) Sample purity distribution across DNMT3A subclasses in Li2016 cohort. Purities were computed using ABSOLUTE. (b) Distribution of the entropy of cell type proportion across DNMT3A subclasses in Li2016 cohort. Cell type proportions were computed using CIBERSORT. The center line denotes the median, the upper and lower box limits denote upper and lower quartiles, and the whiskers denote 1.5× interquartile range.

**Supplementary Figure 8.**
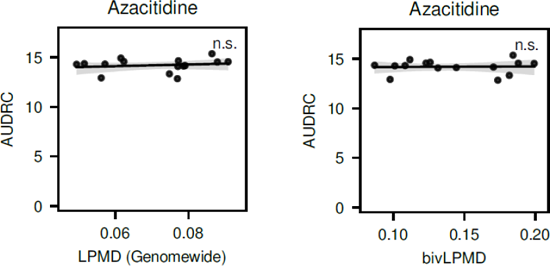
Responses of AML cell lines to azacitidine. Association between genomewide (left) or bivalent-domain-specific LPMD (bivLPMD; right) and area under dose response curve (AUDRC) are shown. No significant correlation was observed (n.s.).

**Supplementary Figure 9.**
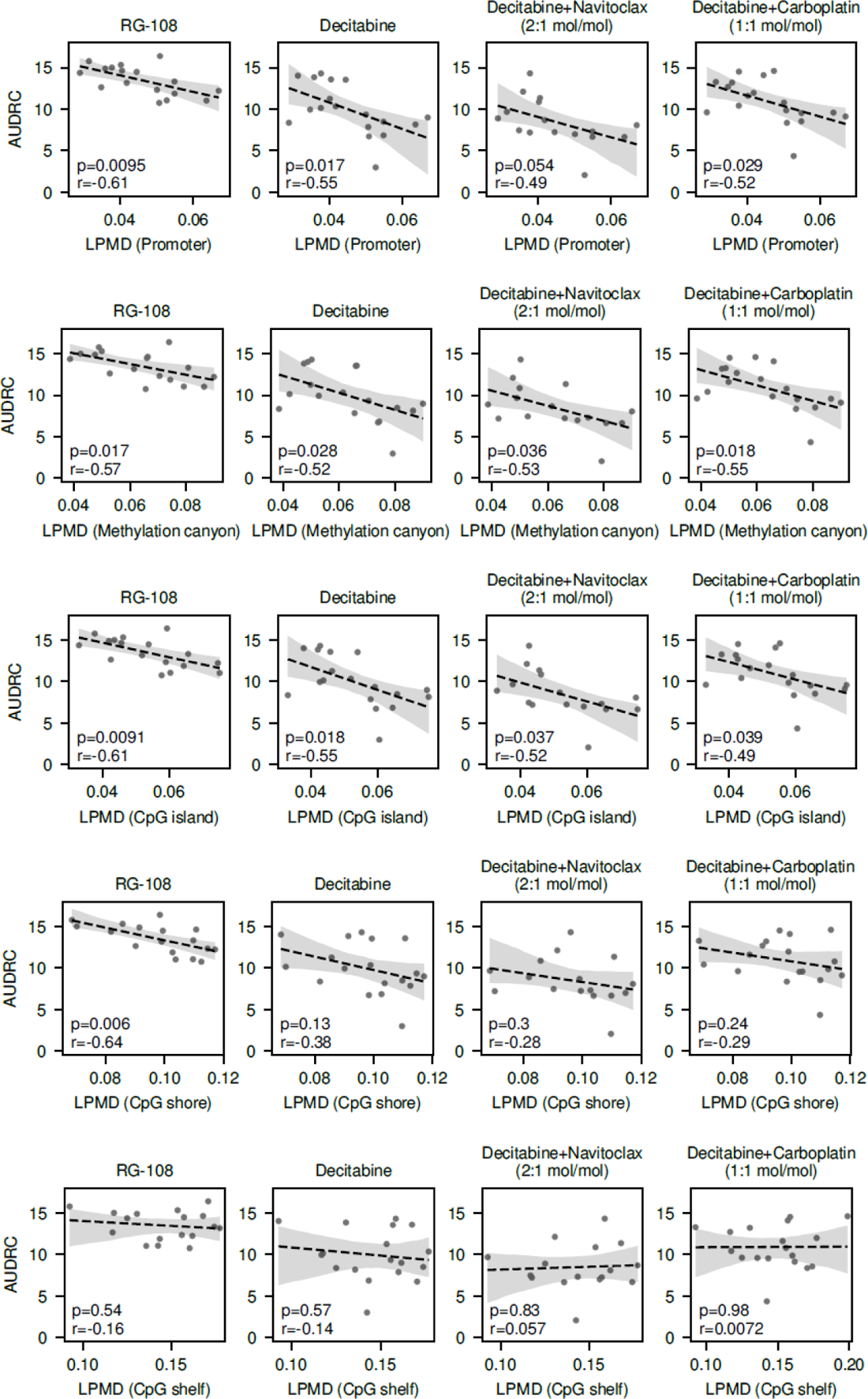
Association between regulatory region-specific LPMD and response to hypomethylating agent. Association between LPMD values within promoters, methylation canyons, CpG islands, CpG shores and CpG shelves and AUDRC for hypomethylating agent treatment are shown. Pearson’s correlation coefficient and corresponding p-values are shown.

**Supplementary Figure 10.**
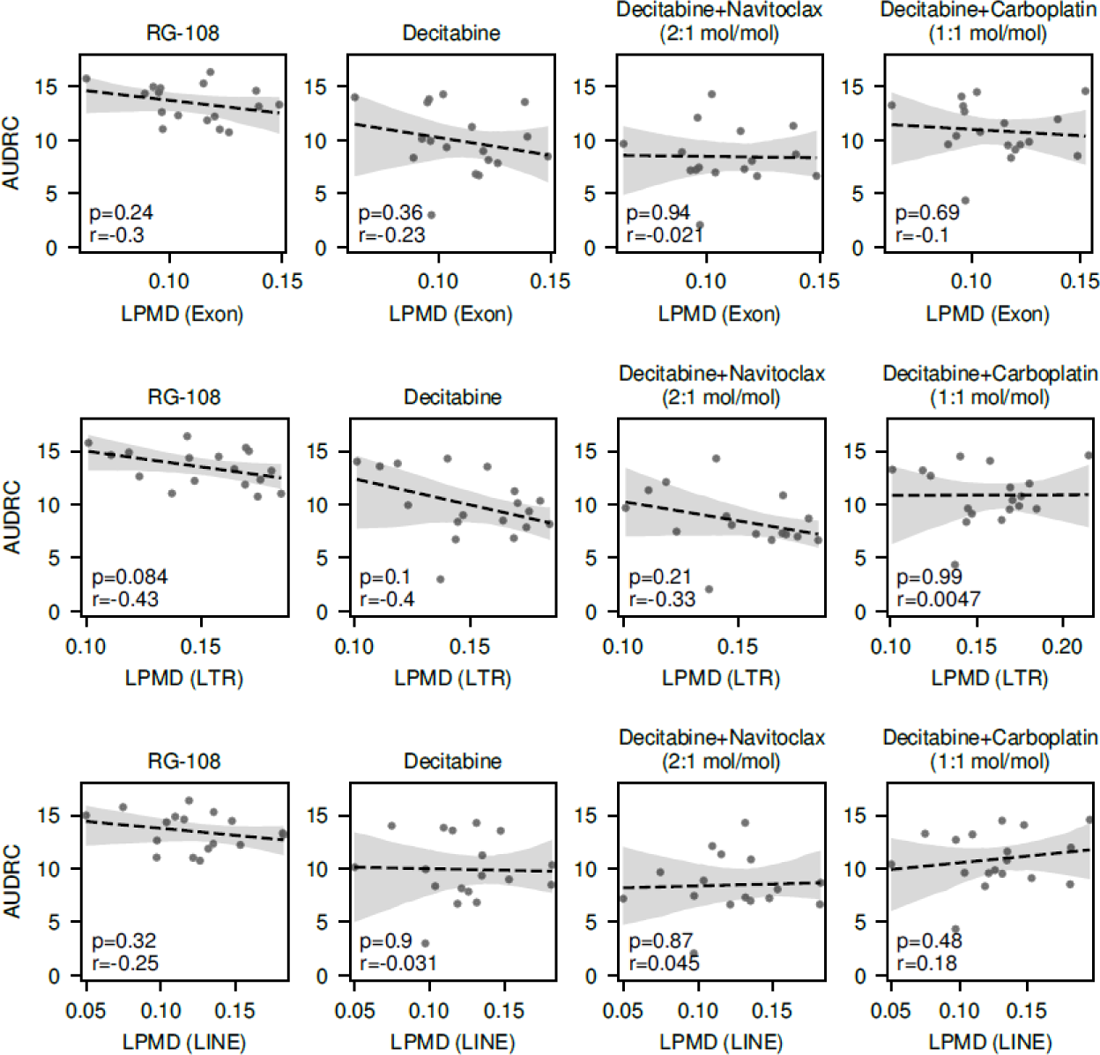
Association between non-regulatory region-specific LPMD and response to hypomethylating agent. Association between LPMD values within exons, LTRs and LINEs and AUDRC for hypomethylating agent treatment are shown. Pearson’s correlation coefficient and corresponding p-values are shown.

**Supplementary Figure 11.**
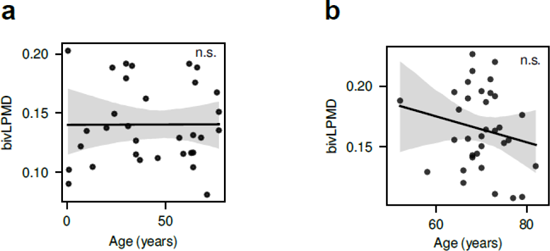
LPMD at bivalent domains are not associated with age. (a) Relationship between cell line age at sampling and bivLPMD values in CCLE AML cell lines. No significant correlation was observed (n.s.). (b) Relationship between patient ages and bivLPMD values in our own cohort for retrospective HMA response analysis (see Figure 5f, g). No significant correlation was observed (n.s.).

**Supplementary Figure 12.**
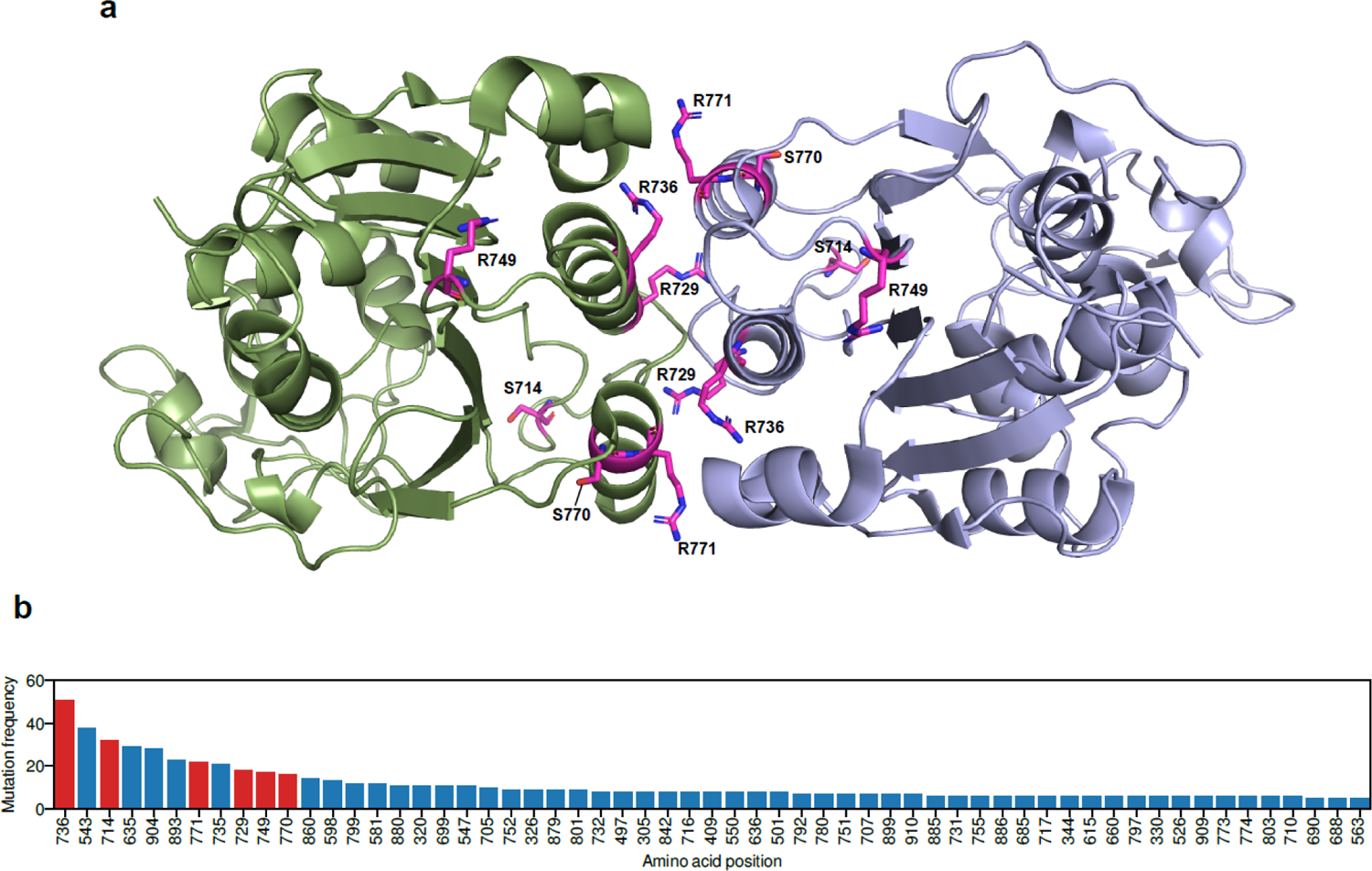
Prevalence of non-R882 mutations that may contribute to dimeric preference of DNMT3A through weakening interactions in tetramer interface. (a) Structure of DNMT3A tetramer interface. Destabilizing residues (stability score < 0.75) that are in the vicinity of the tetramer interface are highlighted. DNMT3A tetramer status was modeled based on DMNT3A-DMNT3L heterotetramer (PDB ID 6BRR) by superimposing DMNT3A on DMNT3L. (b) Top 60 DNMT3A residues that are most frequently mutated in hematological malignancies are shown. Data was downloded from COSMIC. Frequency for R882 (n=1748) was not shown for the visibility. Red bars denote the destabilizing amino acid residues that are placed in the vicinity of tetramer interface of DNMT3A.

## Supplementary Tables

**Supplementary Table 1.** Destabilizing and non-stabilizing residues. Position refers to the 1-based position of an amino acid residue.

| Position | Stability ratio (normalized to WT) | Destabilizing |
| --- | --- | --- |
| 12 | 0.8555979 | No |
| 30 | 0.8360113 | No |
| 90 | 1.084197 | No |
| 181 | 1.044739 | No |
| 183 | 1.060101 | No |
| 192 | 1.047929 | No |
| 254 | 0.4813288 | Yes |
| 267 | 1.016953 | No |
| 272 | 1.022709 | No |
| 281 | 1.017266 | No |
| 288 | 1.017836 | No |
| 290 | 0.4599451 | Yes |
| 292 | 1.014288 | No |
| 293 | 0.3443498 | Yes |
| 295 | 0.3435179 | Yes |
| 297 | 0.1578041 | Yes |
| 298 | 0.3447503 | Yes |
| 299 | 1.024477 | No |
| 301 | 1.012748 | No |
| 302 | 1.032172 | No |
| 304 | 0.9931188 | No |
| 307 | 0.1565844 | Yes |
| 308 | 0.342584 | Yes |
| 309 | 0.5492893 | Yes |
| 310 | 0.1712749 | Yes |
| 312 | 0.6274686 | Yes |
| 314 | 1.072061 | No |
| 316 | 1.071478 | No |
| 318 | 0.9328556 | No |
| 322 | 1.053182 | No |
| 324 | 0.4608576 | Yes |
| 326 | 0.252030067 | Yes |
| 328 | 0.2114281 | Yes |
| 332 | 0.373928 | Yes |
| 337 | 0.3451776 | Yes |
| 339 | 0.1308347 | Yes |
| 342 | 0.8949588 | No |
| 344 | 0.1882434 | Yes |
| 352 | 0.966761 | No |
| 359 | 0.5646341 | Yes |
| 361 | 0.8742617 | No |
| 365 | 0.1697705 | Yes |
| 366 | 0.65244465 | Yes |
| 368 | 0.3986021 | Yes |
| 369 | 0.3906805 | Yes |
| 376 | 0.3382373 | Yes |
| 378 | 1.0956 | No |
| 379 | 0.916074 | No |
| 380 | 0.9477463 | No |
| 385 | 1.072299 | No |
| 400 | 1.15719 | No |
| 407 | 0.6492172 | Yes |
| 410 | 0.2881317 | Yes |
| 413 | 0.8793166 | No |
| 414 | 0.2475144 | Yes |
| 419 | 0.9446362 | No |
| 424 | 0.9331912 | No |
| 426 | 0.9512829 | No |
| 428 | 0.973034 | No |
| 431 | 0.9554172 | No |
| 436 | 0.9696032 | No |
| 437 | 1.043213 | No |
| 438 | 1.000848 | No |
| 455 | 1.150239 | No |
| 458 | 1.0636 | No |
| 474 | 1.130067 | No |
| 477 | 1.098108 | No |
| 478 | 1.088251 | No |
| 484 | 1.138854 | No |
| 494 | 0.4751771 | Yes |
| 495 | 0.9011877 | No |
| 497 | 0.2767733 | Yes |
| 507 | 0.5604438 | Yes |
| 508 | 0.5859341 | Yes |
| 511 | 0.5064597 | Yes |
| 514 | 0.5244856 | Yes |
| 517 | 0.5214359 | Yes |
| 518 | 1.109377 | No |
| 525 | 0.8306941 | No |
| 527 | 0.8820262 | No |
| 529 | 0.8778251 | No |
| 531 | 1.107685 | No |
| 532 | 0.7987965 | No |
| 533 | 0.8857338 | No |
| 537 | 0.4448991 | Yes |
| 543 | 0.9069754 | No |
| 545 | 1.100344 | No |
| 547 | 0.9057181 | No |
| 548 | 0.8853955 | No |
| 549 | 0.2513217 | Yes |
| 550 | 0.848029 | No |
| 554 | 0.579103 | Yes |
| 556 | 1.040666 | No |
| 562 | 0.4254519 | Yes |
| 563 | 0.8841944 | No |
| 567 | 0.8097687 | No |
| 571 | 1.04459 | No |
| 572 | 1.0849 | No |
| 573 | 1.00304 | No |
| 575 | 1.062243 | No |
| 579 | 1.09785 | No |
| 580 | 1.087125 | No |
| 581 | 0.767763 | No |
| 583 | 0.4330766 | Yes |
| 586 | 0.4554169 | Yes |
| 596 | 1.063925 | No |
| 598 | 0.7779466 | No |
| 604 | 0.8633119 | No |
| 623 | 0.7358614 | Yes |
| 627 | 0.8867821 | No |
| 631 | 0.4886155 | Yes |
| 635 | 0.40598 | Yes |
| 636 | 0.3582201 | Yes |
| 637 | 0.2517871 | Yes |
| 638 | 0.4007301 | Yes |
| 639 | 0.3937565 | Yes |
| 641 | 0.7750524 | No |
| 642 | 0.6365951 | Yes |
| 645 | 0.856708 | No |
| 646 | 0.4442819 | Yes |
| 647 | 0.4245519 | Yes |
| 648 | 0.3928059 | Yes |
| 649 | 0.5172071 | Yes |
| 650 | 0.4159871 | Yes |
| 653 | 0.3714397 | Yes |
| 657 | 0.4237124 | Yes |
| 659 | 0.7358019 | Yes |
| 660 | 0.3594055 | Yes |
| 661 | 0.4046589 | Yes |
| 662 | 0.4288399 | Yes |
| 663 | 0.4802697 | Yes |
| 665 | 1.060347 | No |
| 668 | 0.8538262 | No |
| 669 | 0.7318646 | Yes |
| 672 | 0.8466228 | No |
| 675 | 0.9984979 | No |
| 676 | 0.5809479 | Yes |
| 677 | 0.8412493 | No |
| 684 | 0.3898449 | Yes |
| 685 | 0.4403994 | Yes |
| 686 | 0.3386339 | Yes |
| 687 | 0.4086413 | Yes |
| 688 | 0.8012436 | No |
| 690 | 0.3550435 | Yes |
| 691 | 0.4553243 | Yes |
| 695 | 0.9120905 | No |
| 699 | 0.3200215 | Yes |
| 700 | 0.3495642 | Yes |
| 701 | 0.4556982 | Yes |
| 702 | 0.3932404 | Yes |
| 703 | 0.4818903 | Yes |
| 704 | 0.6508228 | Yes |
| 705 | 0.3875358 | Yes |
| 706 | 0.387481 | Yes |
| 707 | 0.3802217 | Yes |
| 709 | 0.8902627 | No |
| 710 | 0.9437781 | No |
| 714 | 0.6880003 | Yes |
| 716 | 1.033337 | No |
| 717 | 0.9261547 | No |
| 718 | 0.7192777 | Yes |
| 720 | 1.061631 | No |
| 728 | 0.8359543 | No |
| 729 | 0.363541 | Yes |
| 731 | 0.40776975 | Yes |
| 733 | 0.426707 | Yes |
| 734 | 0.3905955 | Yes |
| 735 | 0.8630975 | No |
| 736 | 0.3158816 | Yes |
| 737 | 0.4684402 | Yes |
| 741 | 0.6665348 | Yes |
| 742 | 0.3868087 | Yes |
| 743 | 0.4593113 | Yes |
| 747 | 0.7108312 | Yes |
| 749 | 0.3392981 | Yes |
| 751 | 0.3471555 | Yes |
| 752 | 0.3410564 | Yes |
| 754 | 0.3862334 | Yes |
| 755 | 0.4103273 | Yes |
| 756 | 0.9537112 | No |
| 758 | 0.8622715 | No |
| 759 | 0.96031345 | No |
| 761 | 0.9529492 | No |
| 768 | 0.636552 | Yes |
| 769 | 0.4562491 | Yes |
| 770 | 0.4189152 | Yes |
| 771 | 0.5270026 | Yes |
| 772 | 0.7405099 | Yes |
| 774 | 0.2667145 | Yes |
| 777 | 0.4505224 | Yes |
| 778 | 0.3130035 | Yes |
| 780 | 0.4925803 | Yes |
| 781 | 0.9319379 | No |
| 783 | 1.10281 | No |
| 789 | 1.081251 | No |
| 792 | 0.9790729 | No |
| 793 | 0.4067298 | Yes |
| 794 | 0.6260574 | Yes |
| 795 | 0.3420015 | Yes |
| 796 | 0.5078561 | Yes |
| 797 | 0.4046881 | Yes |
| 798 | 0.40725 | Yes |
| 799 | 0.325236 | Yes |
| 800 | 0.9266615 | No |
| 801 | 0.4246202 | Yes |
| 803 | 0.9229618 | No |
| 804 | 1.028859 | No |
| 811 | 0.9130292 | No |
| 813 | 1.025642 | No |
| 814 | 1.049149 | No |
| 815 | 0.9383628 | No |
| 822 | 1.007593 | No |
| 825 | 0.9238034 | No |
| 826 | 1.062891 | No |
| 828 | 1.062804 | No |
| 829 | 0.9500532 | No |
| 835 | 0.9569821 | No |
| 836 | 0.9760696 | No |
| 838 | 0.9859692 | No |
| 849 | 0.5965496 | Yes |
| 850 | 1.02501 | No |
| 857 | 0.937554 | No |
| 860 | 0.9209173 | No |
| 865 | 0.5922805 | Yes |
| 868 | 0.9252796 | No |
| 869 | 0.5570258 | Yes |
| 872 | 1.025123 | No |
| 873 | 1.041027 | No |
| 879 | 0.8984118 | No |
| 880 | 0.9616624 | No |
| 881 | 0.6161368 | Yes |
| 882 | 0.9764945 | No |
| 884 | 1.002667 | No |
| 886 | 1.072883 | No |
| 893 | 0.9518706 | No |
| 896 | 0.8088213 | No |
| 898 | 0.7921419 | No |
| 899 | 0.8239997 | No |
| 901 | 0.4658968 | Yes |
| 902 | 0.36474025 | Yes |
| 904 | 0.2603852 | Yes |
| 905 | 0.4671688 | Yes |
| 906 | 0.6004309 | Yes |
| 907 | 0.3925456 | Yes |
| 908 | 0.4078807 | Yes |
| 909 | 0.327513 | Yes |
| 910 | 0.5581608 | Yes |
| 911 | 0.5075343 | Yes |

**Supplementary Table 2.** Characteristics of SNUH AML patients for bisulfite sequencing analysis.

| Sample | Age (yr) | Sex | Karyotype |
| --- | --- | --- | --- |
| SNUH4794 | 46 | F | 46,XX,t(8;21)(q22;q22)[20] |
| SNUH5053 | 48 | M | 45,X,-Y,t(8;21)(q22;q22)[20] |
| SNUH5070 | 51 | F | 46,XX[20] |
| SNUH5160 | 41 | M | 46,XY[20] |
| SNUH5174 | 56 | F | Unknown |
| SNUH5196 | 64 | F | 46,XX,del(13)(q12q14)[16]/46,XX[5] |
| SNUH5323 | 43 | M | 46,XY[20] |
| SNUH5347 | 64 | M | 44,XY,-3,add(5)(q?15),der(7)t(7;12)(p13;q13)add(7)(q22),+8,-10,-12,add(17)(q21)[17]/88,slx2[1]/46,XY[2] |
| SNUH5553 | 68 | M | 46,XY,der(7)t(7;14)(q22;q24),add(18)(q21.1)[21] |
| SNUH5576 | 77 | F | 47,XX,+4[20] |
| SNUH5696 | 66 | M | 46,XY[21] |
| SNUH5763 | 71 | M | 47,XY,+10[4]/46,XY[17] |
| SNUH6002 | 54 | F | 46,X,t(8;21)(q22;q22)[18]/46,XX[2] |
| SNUH6076 | 79 | F | 46,XX[11] |
| SNUH6407 | 65 | F | 46,XX,t(8;21)(q22;q22)[17]/46,XX[4] |
| SNUH4939 | 68 | F | 46,XX[20] |
| SNUH5253 | 61 | F | 46,XX[20] |
| SNUH5996 | 61 | M | 46,XY[20] |
| SNUH5997 | 47 | M | 44,XY,der(3;5)(q10;p10),add(7)(p22),-8,-9,add(17)(p13),+r,~100dmin[19]/88,idemx2[1].nuc ish(MYC amp)[192/200] |
| SNUH6446 | 55 | F | 46,XX,add(12)(q12)[12]/46,idem,add(7)(p22)[2]/46,XX[6] |

**Supplementary Table 3.** Characteristics of patients with hematological disorders for HMA response analysis.

| **Sample** | **Age (yr)** | **Sex** | **Disease classification** |
| --- | --- | --- | --- |
| SNUH2715 | 68 | F | Acute Myeloid Leukemia |
| SNUH2789 | 79 | F | Acute Myeloid Leukemia |
| SNUH2996 | 68 | M | Acute Myeloid Leukemia |
| SNUH3173 | 73 | M | Acute Myeloid Leukemia |
| SNUH3190 | 77 | F | Acute Myeloid Leukemia |
| SNUH3313 | 67 | F | Acute Myeloid Leukemia |
| SNUH3626 | 73 | M | Acute Myeloid Leukemia |
| SNUH3651 | 70 | M | Acute Myeloid Leukemia |
| SNUH4028 | 73 | M | Acute Myeloid Leukemia |
| SNUH4390 | 82 | M | Acute Myeloid Leukemia |
| SNUH4590 | 74 | F | Acute Myeloid Leukemia |
| SNUH4819 | 70 | F | Acute Myeloid Leukemia |
| SNUH4871 | 72 | M | Acute Myeloid Leukemia |
| SNUH4995 | 70 | F | Acute Myeloid Leukemia |
| SNUH5018 | 68 | M | Acute Myeloid Leukemia |
| SNUH5137 | 64 | M | Acute Myeloid Leukemia |
| SNUH5167 | 68 | M | Acute Myeloid Leukemia |
| SNUH5189 | 66 | M | Acute Myeloid Leukemia |
| SNUH5199 | 76 | F | Acute Myeloid Leukemia |
| SNUH7090 | 75 | F | Acute Myeloid Leukemia |
| SNUH7098 | 58 | F | Myelodysplastic Syndrome (RAEB) |
| SNUH7103 | 71 | F | Acute Myeloid Leukemia |
| SNUH7196 | 67 | M | Acute Myeloid Leukemia |
| SNUH7247 | 66 | F | Myelodysplastic Syndrome (t-MDS) |
| SNUH7349 | 70 | M | Acute Myeloid Leukemia |
| SNUH7366 | 73 | M | Acute Myeloid Leukemia |
| SNUH7383 | 67 | F | Myelodysplastic Syndrome (t-MDS) |
| SNUH7509 | 69 | M | Acute Myeloid Leukemia |
| SNUH7549 | 52 | F | Myelodysplastic Syndrome (RAEB) |
| SNUH7666 | 71 | M | Myelodysplastic Syndrome (RAEB) |
| SNUH7683 | 72 | M | Acute Myeloid Leukemia |
| SNUH7692 | 65 | M | Myelodysplastic Syndrome (RAEB) |
| SNUH7720 | 64 | M | Myelodysplastic Syndrome (RAEB) |
| SNUH7783 | 79 | M | Acute Myeloid Leukemia |

## Notes

### Competing Interest Statement

Y. Koh. is the founder and CEO of Genome Opinion. C. Sun. is the executive director and owns stock in Genome Opinion. The other authors declare no competing interests.

## References

1. Yang L., Rau R., Goodell M. A. DNMT3A in haematological malignancies. Nat Rev Cancer 15, 152–165 (2015).

2. Chen T., Ueda Y., Dodge J. E., Wang Z., Li E. Establishment and maintenance of genomic methylation patterns in mouse embryonic stem cells by Dnmt3a and Dnmt3b. Mol Cell Biol 23, 5594–5605 (2003).

3. Brunetti L., Gundry M. C., Goodell M. A. DNMT3A in Leukemia. Cold Spring Harb Perspect Med 7, (2017).

4. Russler-Germain D. A., et al. The R882H DNMT3A mutation associated with AML dominantly inhibits wild-type DNMT3A by blocking its ability to form active tetramers. Cancer Cell 25, 442–454 (2014).

5. Emperle M., et al. The DNMT3A R882H mutation does not cause dominant negative effects in purified mixed DNMT3A/R882H complexes. Sci Rep 8, 13242 (2018).

6. Huang Y. H., et al. Systematic Profiling of DNMT3A Variants Reveals Protein Instability Mediated by the DCAF8 E3 Ubiquitin Ligase Adaptor. Cancer Discov, (2021).

7. Landau D. A., et al. Locally disordered methylation forms the basis of intratumor methylome variation in chronic lymphocytic leukemia. Cancer Cell 26, 813–825 (2014).

8. Li S., et al. Distinct evolution and dynamics of epigenetic and genetic heterogeneity in acute myeloid leukemia. Nat Med 22, 792–799 (2016).

9. Sheffield N. C., et al. DNA methylation heterogeneity defines a disease spectrum in Ewing sarcoma. Nat Med 23, 386–395 (2017).

10. Pan H., et al. Epigenomic evolution in diffuse large B-cell lymphomas. Nat Commun 6, 6921 (2015).

11. Li S., et al. Somatic Mutations Drive Specific, but Reversible, Epigenetic Heterogeneity States in AML. Cancer Discov 10, 1934–1949 (2020).

12. Cancer Genome Atlas Research N., et al. Genomic and epigenomic landscapes of adult de novo acute myeloid leukemia. N Engl J Med 368, 2059–2074 (2013).

13. Varadi M., et al. AlphaFold Protein Structure Database: massively expanding the structural coverage of protein-sequence space with high-accuracy models. Nucleic Acids Res 50, D439–D444 (2022).

14. Holz-Schietinger C., Matje D. M., Harrison M. F., Reich N. O. Oligomerization of DNMT3A controls the mechanism of de novo DNA methylation. J Biol Chem 286, 41479–41488 (2011).

15. Holz-Schietinger C., Matje D. M., Reich N. O. Mutations in DNA methyltransferase (DNMT3A) observed in acute myeloid leukemia patients disrupt processive methylation. J Biol Chem 287, 30941–30951 (2012).

16. Lee D., Koo B., Yang J., Kim S. Metheor: Ultrafast DNA methylation heterogeneity calculation from bisulfite read alignments. Preprint at https://www.biorxiv.org/content/10.1101/2022.07.20.500893v1 (2022).

17. Spencer D. H., et al. CpG Island Hypermethylation Mediated by DNMT3A Is a Consequence of AML Progression. Cell 168, 801–816 e813 (2017).

18. Juhling F., Kretzmer H., Bernhart S. H., Otto C., Stadler P. F., Hoffmann S. metilene: fast and sensitive calling of differentially methylated regions from bisulfite sequencing data. Genome Res 26, 256–262 (2016).

19. Encode Project Consortium. An integrated encyclopedia of DNA elements in the human genome. Nature 489, 57–74 (2012).

20. Ernst J., Kellis M. Chromatin-state discovery and genome annotation with ChromHMM. Nat Protoc 12, 2478–2492 (2017).

21. Scherer M., et al. Quantitative comparison of within-sample heterogeneity scores for DNA methylation data. Nucleic Acids Res 48, e46 (2020).

22. Landan G., et al. Epigenetic polymorphism and the stochastic formation of differentially methylated regions in normal and cancerous tissues. Nature Genetics 44, 1207–1214 (2012).

23. Bernstein B. E., et al. A bivalent chromatin structure marks key developmental genes in embryonic stem cells. Cell 125, 315–326 (2006).

24. Fleming H. E., et al. Wnt signaling in the niche enforces hematopoietic stem cell quiescence and is necessary to preserve self-renewal in vivo. Cell Stem Cell 2, 274–283 (2008).

25. Huang J., Nguyen-McCarty M., Hexner E. O., Danet-Desnoyers G., Klein P. S. Maintenance of hematopoietic stem cells through regulation of Wnt and mTOR pathways. Nat Med 18, 1778–1785 (2012).

26. Wainwright E. N., Scaffidi P. Epigenetics and Cancer Stem Cells: Unleashing, Hijacking, and Restricting Cellular Plasticity. Trends Cancer 3, 372–386 (2017).

27. Santini V., Ossenkoppele G. J. Hypomethylating agents in the treatment of acute myeloid leukemia: A guide to optimal use. Crit Rev Oncol Hematol 140, 1–7 (2019).

28. Kordella C., Lamprianidou E., Kotsianidis I. Mechanisms of Action of Hypomethylating Agents: Endogenous Retroelements at the Epicenter. Front Oncol 11, 650473 (2021).

29. Sigalotti L., et al. Epigenetic drugs as pleiotropic agents in cancer treatment: biomolecular aspects and clinical applications. J Cell Physiol 212, 330–344 (2007).

30. Agrawal K., Das V., Vyas P., Hajduch M. Nucleosidic DNA demethylating epigenetic drugs - A comprehensive review from discovery to clinic. Pharmacol Ther 188, 45–79 (2018).

31. Tyner J. W., et al. Functional genomic landscape of acute myeloid leukaemia. Nature 562, 526–531 (2018).

32. Glass J. L., et al. Epigenetic Identity in AML Depends on Disruption of Nonpromoter Regulatory Elements and Is Affected by Antagonistic Effects of Mutations in Epigenetic Modifiers. Cancer Discov 7, 868–883 (2017).

33. Sandoval J. E., Huang Y. H., Muise A., Goodell M. A., Reich N. O. Mutations in the DNMT3A DNA methyltransferase in acute myeloid leukemia patients cause both loss and gain of function and differential regulation by protein partners. J Biol Chem 294, 4898–4910 (2019).

34. Martin M. Cutadapt removes adapter sequences from high-throughput sequencing reads. EMBnet journal 17, 10–12 (2011).

35. Krueger F., Andrews S. R. Bismark: a flexible aligner and methylation caller for Bisulfite-Seq applications. Bioinformatics 27, 1571–1572 (2011).

36. Ryan D. MethylDackel. https://github.com/dpryan79/MethylDackel (2022).

37. Cole C. B., et al. Haploinsufficiency for DNA methyltransferase 3A predisposes hematopoietic cells to myeloid malignancies. J Clin Invest 127, 3657–3674 (2017).

38. Li H., Durbin R. Fast and accurate short read alignment with Burrows-Wheeler transform. Bioinformatics 25, 1754–1760 (2009).

39. Kim S., et al. Strelka2: fast and accurate calling of germline and somatic variants. Nat Methods 15, 591–594 (2018).

40. Koboldt D. C., et al. VarScan: variant detection in massively parallel sequencing of individual and pooled samples. Bioinformatics 25, 2283–2285 (2009).

41. Cingolani P., et al. A program for annotating and predicting the effects of single nucleotide polymorphisms, SnpEff: SNPs in the genome of Drosophila melanogaster strain w1118; iso-2; iso-3. Fly (Austin) 6, 80-92 (2012).

42. Cingolani P., et al. Using Drosophila melanogaster as a Model for Genotoxic Chemical Mutational Studies with a New Program, SnpSift. Front Genet 3, 35 (2012).

43. Cerami E., et al. The cBio cancer genomics portal: an open platform for exploring multidimensional cancer genomics data. Cancer Discov 2, 401–404 (2012).

44. Vaisvila R., et al. Enzymatic methyl sequencing detects DNA methylation at single-base resolution from picograms of DNA. Genome Res 31, 1280–1289 (2021).

45. Quinlan A. R., Hall I. M. BEDTools: a flexible suite of utilities for comparing genomic features. Bioinformatics 26, 841–842 (2010).

46. Frankish A., et al. Gencode 2021. Nucleic Acids Res 49, D916–D923 (2021).

47. Jeong M., et al. Large conserved domains of low DNA methylation maintained by Dnmt3a. Nat Genet 46, 17–23 (2014).

48. Ghandi M., et al. Next-generation characterization of the Cancer Cell Line Encyclopedia. Nature 569, 503–508 (2019).

49. Amemiya H. M., Kundaje A., Boyle A. P. The ENCODE Blacklist: Identification of Problematic Regions of the Genome. Sci Rep 9, 9354 (2019).

50. Rees M. G., et al. Correlating chemical sensitivity and basal gene expression reveals mechanism of action. Nat Chem Biol 12, 109–116 (2016).

51. Gadagkar S. R., Call G. B. Computational tools for fitting the Hill equation to dose-response curves. J Pharmacol Toxicol Methods 71, 68–76 (2015).

